# InfluenceNet: Encoding Motif Influence for Interpretable Modeling of Cis-Regulatory Syntax

**DOI:** 10.1101/2025.11.20.689553

**Authors:** Jieni Hu, Julianne Fetter, David R. Koes, Maria Chikina

**Affiliations:** Joint Carnegie Mellon-University of Pittsburgh Ph.D. Program in Computational Biology, Pittsburgh, PA, USA; Department of Computational and Systems Biology, University of Pittsburgh, Pittsburgh, PA, USA; Saint Vincent College, Latrobe, PA, USA

**Keywords:** Machine learning, Cis-regulatory grammar, Interpretation

## Abstract

Cis-regulatory elements shape gene expression by recruiting transcription factors (TFs) to DNA motifs, yet how motifs cooperate or compete across distances remains poorly understood. Existing deep learning models predict TF binding with high accuracy but rely on computationally intensive post-hoc analyses to infer regulatory syntax, limiting interpretability and scalability. Here we present InfluenceNet, a transparent convolutional architecture that directly encodes motif influence-a position– and motif-specific profile quantifying how one motif modulates binding at another. Trained on ChIP-nexus data for pluripotency TFs, InfluenceNet achieves performance comparable to state-of-the-art models while providing immediate, global interpretability. The learned influence profiles recapitulate known principles of motif syntax, including Oct4-Sox2 pioneering activity and Nanog’s striking 10.5-bp helical periodicity, and reveal a cooperative Nanog-Nanog grammar supported by de novo motif discovery. Integrating these results with structural and molecular-dynamics modeling, we propose that Nanog forms chain-like oligomers on DNA through asymmetric contacts between WRD (tryptophan repeat domain) and DBD (DNA-binding domain) that maintain helical phase alignment. Together, these findings establish InfluenceNet as a general framework for accurate prediction and mechanistic interpretation of cis-regulatory grammar, bridging deep learning and biophysical insight.

## 1 Introduction

Cis-regulatory elements (CREs) modulate gene expression by recruiting transcription factors (TFs) to specific DNA motifs within accessible chromatin regions. While the presence of motifs is necessary, it is far from sufficient to explain TF binding. Binding is strongly modulated by the spatial arrangement and combination of motifs, a phenomenon often described as cis-regulatory grammar. Deciphering this grammar is essential for predicting the impact of noncoding variants on CRE function and, ultimately, on gene regulation.

Two distinct views of cis-regulatory grammar have emerged. In some loci, binding follows hard grammar rules, where precise motif identities, spacing, and orientations are required. The interferon-stimulated response element (ISRE), for example, encodes strict pairwise positioning of IRF and STAT motifs to ensure cooperative assembly of the enhanceosome. Such cases demonstrate that rigid syntax can enforce highly specific transcriptional responses. By contrast, most regulatory regions exhibit soft grammar, in which motif interactions are probabilistic and distance-dependent. Here, cooperative or competitive effects decay gradually with spacing and vary across contexts, producing flexible yet reproducible patterns of regulation. While hard grammar is mechanistically compelling in isolated examples, soft grammar appears to dominate genome-wide enhancer logic.

In recent years, high-throughput genomics has provided increasingly detailed functional readouts of CRE activity. Chromatin immunoprecipitation sequencing (ChIP-seq) remains a standard approach for profiling genome-wide TF binding, while its high-resolution extensions, ChIP-exo [1] and ChIP-nexus [2], achieve nucleotide-resolution mapping that is necessary for dissecting fine-scale combinatorial effects [1, 2]. However, even with such precise measurements, the underlying rules of motif interactions cannot be directly inferred from experimental profiles. Inferring grammar requires computational modeling.

One modeling strategy has been to train high-capacity sequence-to-function (S2F) models that achieve high fidelity in predicting experimental data and then probe them with in silico experiments. These models take DNA sequence as input and predict functional readouts such as TF binding [3], chromatin accessibility [4, 5], transcription initiation [6, 7, 8], 3D genome architecture [9], and gene expression [10, 11]. BPNet [3] exemplifies this approach: it predicts base-resolution ChIP-nexus profiles and, through extensive perturbation analyses, revealed soft motif syntax. In particular, BPNet showed that pluripotency factors display distance-dependent cooperativity and uncovered striking 10.5 bp helical phasing of Nanog binding. These findings underscored the importance of modeling soft grammar explicitly.

However, in BPNet and related architectures, grammar is not encoded directly in the model. Instead, syntax must be reconstructed post hoc through synthetic motif embeddings, mutagenesis, or ablation experiments [3, 5, 7, 8]. Such workflows are powerful but computationally expensive, input-dependent, and difficult to generalize, limiting their ability to provide global or scalable insights. Other approaches, such as attribution methods [12] or rule-based motif effect models like Puffin [6], focus on individual motif contributions but do not directly characterize distance-dependent motif-motif interactions. Overall, despite promising progress, current pipelines leave us without a modeling framework that transparently encodes soft grammar while retaining predictive accuracy.

Here we introduce InfluenceNet, a lightweight convolutional neural network that bridges this gap by directly modeling motif influence: a position– and motif-specific profile quantifying how the presence of one motif enhances or represses binding at another across distances. InfluenceNet is explicitly structured to map onto biological processes—motif detection, motif influence, and footprint generation—providing immediate interpretability without post-hoc analysis.

Applied to ChIP-nexus data for pluripotency TFs, InfluenceNet achieved predictive performance comparable to the state-of-the-art model BPNet while yielding direct mechanistic insight. Its learned influence profiles recapitulated known principles, including Oct4-Sox2 pioneering activity and the Nanog’s helical cooperativity, and further revealed a cooperative Nanog-Nanog grammar via de novo motif discovery. Integrating these findings with structural modeling and molecular dynamics simulations, we propose that Nanog forms chain-like oligomers on DNA through asymmetric WR-DBD contacts that preserve helical phase alignment. More broadly, InfluenceNet establishes a general framework for accurate and interpretable modeling of cis-regulatory grammar across diverse genomic readouts.

## 2 Results

### 2.1 Interpretable model for predicting base-resolution TF binding profiles

While transcription factors preferentially bind to specific DNA motifs, the presence of motifs alone is insufficient to pinpoint binding locations. Binding outcomes depend strongly on combinatorial effects—motifs acting cooperatively or competitively in distance– and orientation-dependent ways. To capture these dependencies, we developed InfluenceNet, a lightweight neural network that encodes motif influence as a one-dimensional positional profile. This explicit representation allows the model to quantify how motifs modulate each other’s binding activity and thereby improve prediction of TF binding events.

InfluenceNet is structured around three convolutional modules (Fig. 1a). The sequence layer detects motif patterns in the input DNA, generating raw motif-driven activations based purely on local sequence. The influence layer computes position– and motif-specific weights that represent cooperative effects across distances; these weights are applied multiplicatively to the raw activations to yield context-aware activations. Finally, the assay layer serves as a readout module that maps these activations to nucleotide-resolution ChIP-nexus profiles, ensuring that the model output aligns with the experimental measurement. Thus, while the first two layers correspond to regulatory processes of motif detection and influence, the assay layer provides a technical bridge to the assay data.

**Figure 1:**
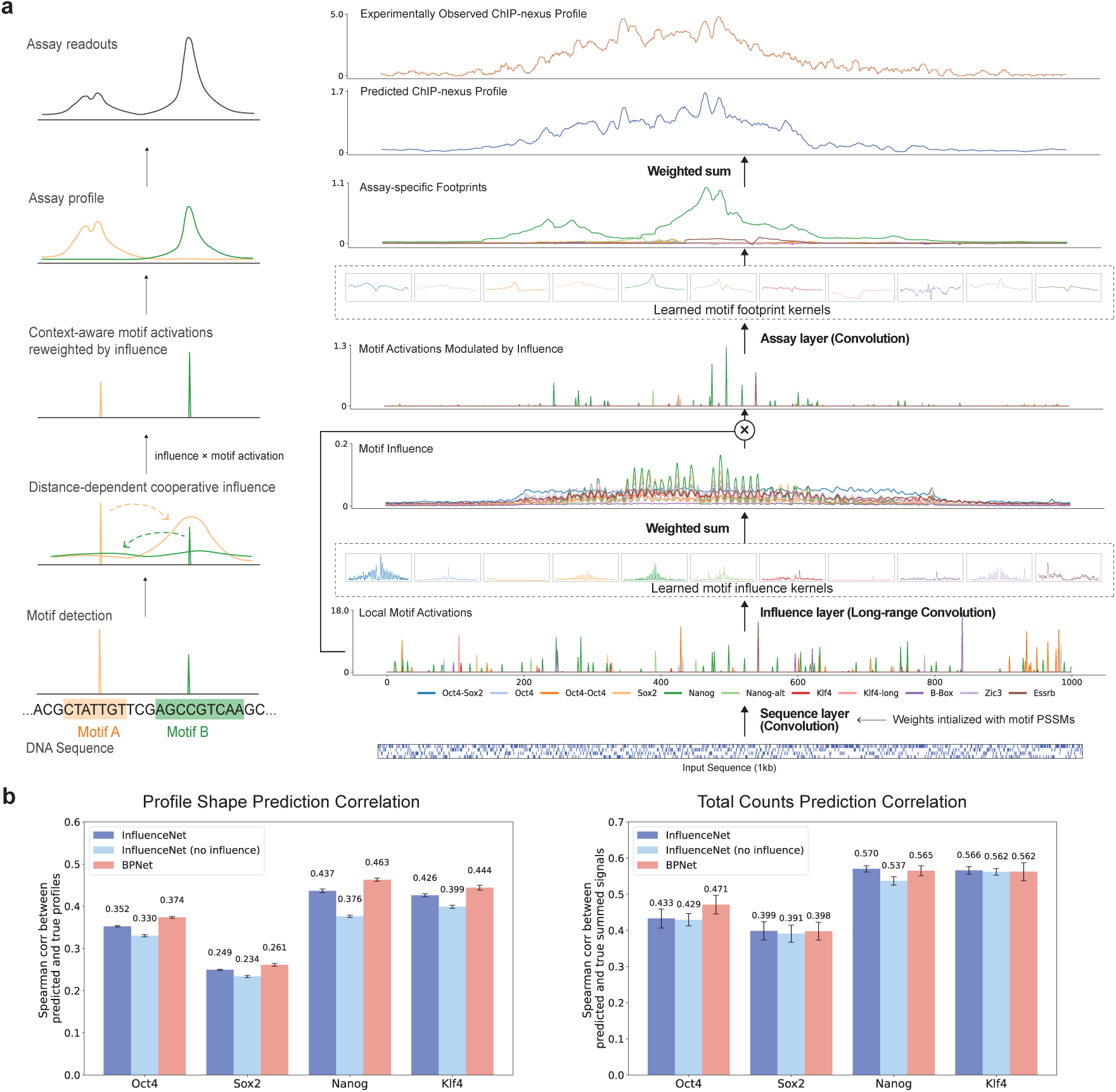
InfluenceNet predicts ChIP-nexus profile from DNA sequence. **a**, Schematic illustration of the InfluenceNet architecture. The model consists of three convolutional modules: a sequence layer, an influence layer, and an assay layer. The sequence layer computes motif activations from local DNA sequence. The influence layer generates motif-specific and distance-dependent influence weights, which are then linearly transformed and applied multiplicatively to the raw activations, yielding influence-modified activations that incorporate broader context. Finally, the assay layer maps the influence-modified activations to predicted TF binding footprints. **b**, Cross-validation performance of InfluenceNet compared to baselines. Performance was evaluated for both profile shape (left) and total counts (right).

We trained InfluenceNet on base-resolution ChIP-nexus data for the pluripotency TFs Oct4, Sox2, Nanog, and Klf4 [3], using position weight matrices (PWMs) of the 11 representative motifs from BPNet to initialize most sequence kernels and leaving four kernels free for de novo motif discovery. We performed chromosome-based fivefold cross-validation and reported average model performance across folds (Fig. 1b). Despite its compact size (∼20k parameters), InfluenceNet achieved accuracy close to BPNet (∼124k parameters). Across TFs, profile correlations for InfluenceNet trailed BPNet by only 1.2-2.6% (Fig. 1b, left), while cross-region correlations matched or exceeded BPNet for all but Oct4 (Fig. 1b, right). These results highlight that a shallow, biologically structured model can perform competitively with a deep architecture while remaining interpretable. Comparing InfluenceNet and BPNet in more details derived similar conclusions, and revealed that the performance across regions was highly correlated with the total number of counts being a large predictor of performance (Supplementary Fig. 1).

To pinpoint the role of the influence layer, we compared InfluenceNet to a variant in which this layer was removed (“InfluenceNet (no influence)”). Eliminating the influence layer substantially reduced accuracy for base-resolution profile prediction, underscoring the importance of explicitly modeling motif interactions for capturing fine-scale mechanistic features of TF binding. By contrast, inter-region correlations were only modestly affected (Fig. 1b). This divergence likely reflects the underlying biology: variation in TF binding across regions carries functional consequences and has been shaped by evolutionary redundancy, whereas the precise shape of local footprints may be less constrained by function and instead provide a window into mechanistic details of TF-motif organization. More broadly, this highlights the value of base-pair resolution data, which has become increasingly central in recent sequence-to-function models [6]. Our results suggest that even complex CNN architectures, which can in principle learn comprehensive motif interactions, may fail to recover meaningful representations when trained on scalar readouts rather than full profiles.

We further tested architectural choices of InfluenceNet for two key parameters: number of free kernels and width of influence layer kernel. Increasing the number of randomly initialized “free kernels” in the first convolution layer improved performance slightly (Supplementary Fig. 3a), indicating that learning additional motifs complements the predefined motifs. Similarly, increasing the width of the influence kernel enhanced predictions, suggesting the benefits of capturing long-range motif dependencies (Supplementary Fig. 3b).

### 2.2 Interpretable model structure mirrors biological modularity

InfluenceNet is explicitly factorized into biologically interpretable modules corresponding to motif detection, motif influence, and assay readout. To verify that each component behaves as intended, we first examined the assay-layer kernels, which map motif activations to experimental footprints. Averaging results from 50 models trained with different random seeds and data splits, InfluenceNet reproduced strong, motif-specific footprint signals for all four TFs (Fig. 2a). The learned footprint kernels closely matched the averaged ChIP-nexus footprints (Supplementary Fig. 4a), confirming that the model accurately captured marginal footprint shapes. Having validated this component, we next focused on the influence layer, which encodes the biologically most informative parameters.

**Figure 2:**
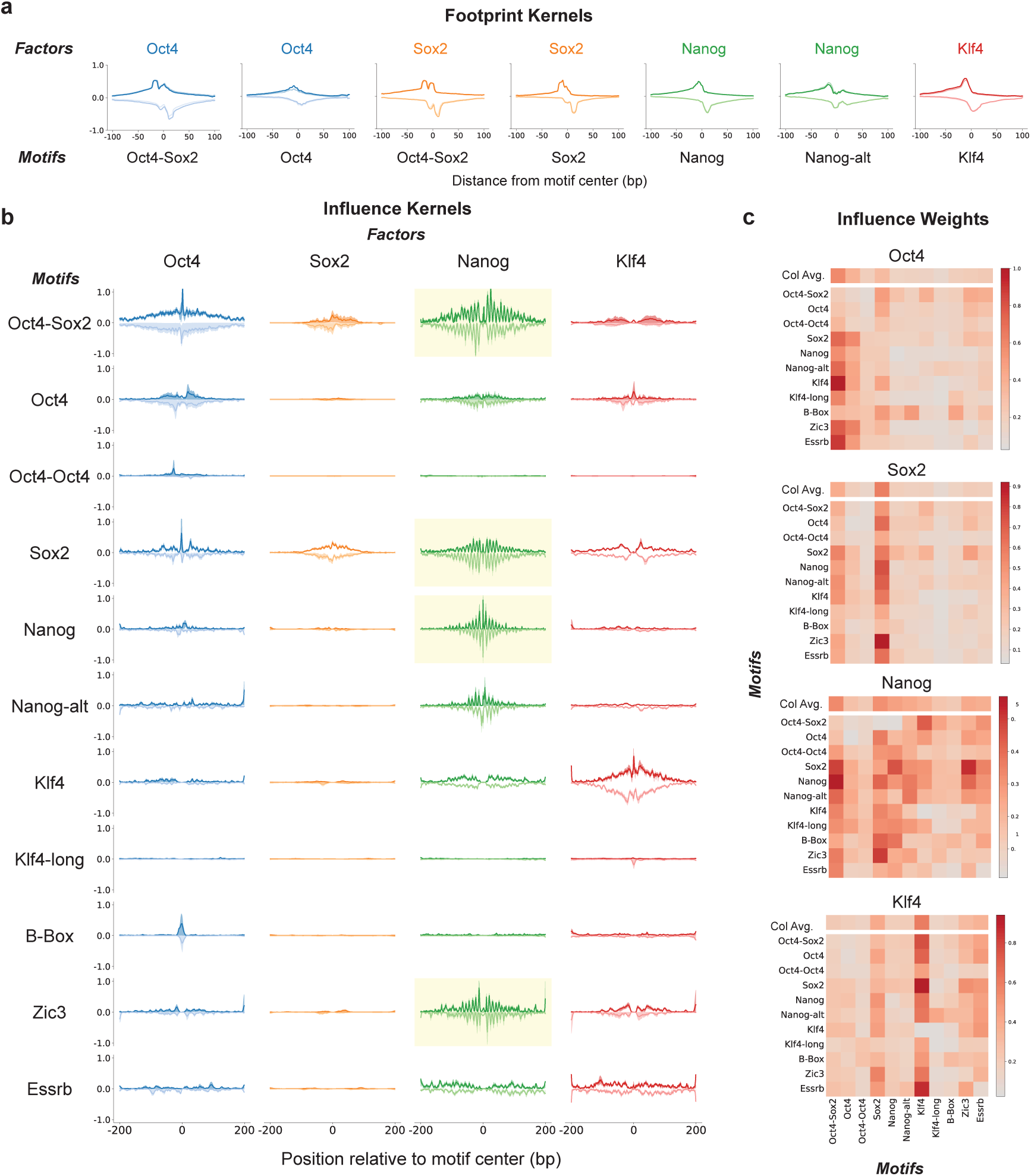
InfluenceNet’s interpretable modules reveal biologically consistent motif syntax. **a**, Assay-layer kernels learned by InfluenceNet accurately recovered representative ChIP-nexus footprints for specific motifs. **b**, Ribbon plots showing the averaged influence kernel weights for each TF, with shaded areas indicating interquartile ranges. Dark and light colors represent influence from the forward and reverse strand motifs, respectively. Motifs that show specifically strong 10.5 ± 0.3 bp periodic influence are highlighted with a light yellow background. **c**, Motif influence weight matrices, extracted from weights of the linear layer that combines influences in weighted sums. The column averages highlight motifs with dominant influence on each TF. All the results represent averages across 50 models (5 cross-validation folds × 10 random seeds) per TF.

InfluenceNet directly learns base-resolution motif-to-motif influence profiles that quantify how the presence of one motif modulates nearby TF binding, either enhancing or repressing occupancy. These spatial influence patterns capture soft motif syntax, the distance-dependent grammar underlying combinatorial regulation. Such syntax is particularly relevant in pluripotent stem cells [13], where Oct4, Sox2, Nanog, and Klf4 frequently co-occupy enhancers [14, 15]. To explore these syntactic rules, we visualized the averaged influence kernels learned by InfluenceNet (Fig. 2b). Each column shows the influence of motifs on a given TF’s binding, averaged over 50 independent models, with shaded areas indicating interquartile variability. Across all motifs, we observed a consistent pattern of decay, that is strongest near motif centers and tapers with distance, indicating locally cooperative yet spatially constrained effects.

Oct4 and Sox2 binding were both dominated by the Oct4-Sox2 composite motif, with Oct4 showing stronger dependence than either individual motif alone. This pattern agrees with their known non-additive cooperative binding [16] and physical interaction [17]. The observed influence extended beyond typical protein-protein contact ranges, suggesting that Oct4/Sox2 may promote neighboring binding through local chromatin opening [18]. Sox2 binding was less influenced by other motifs than Oct4, consistent with its stronger pioneering role [19]. Nanog binding was strongly enhanced by Oct4-Sox2 and Sox2 motifs but exerted little reciprocal influence, revealing directional cooperation that is consistent with Oct4/Sox2’s pioneering activity [20]. The long-range influence from Oct4-Sox2 to Nanog again exceeded direct interaction distances, supporting a model in which Oct4/Sox2 create an open chromatin environment that facilitates Nanog recruitment. Klf4 binding was largely self-dependent, but with modest contributions from Sox2 and Essrb motifs, in line with their previously reported cooperativity [21].

The linear coefficients that reweight motif influences (Fig. 2c) mirrored these trends: Oct4-Sox2 and Sox2 motifs showed the largest column averages for Oct4, Sox2, and Nanog binding, emphasizing their dominant role in shaping binding probabilities, whereas Nanog and Klf4 contributed minimally, consistent with their weaker pioneering activity.

Finally, our framework reveals that the soft syntax governing Nanog binding exhibits a clear 10.5 bp helical periodicity (Fig. 2b), a hallmark previously inferred only through in silico motif perturbations in BPNet [3]. In contrast, InfluenceNet captures this periodic pattern directly and transparently through its learned parameters, without requiring post-hoc experiments. Because each curve represents an average across 50 independently trained models, the recurrence of this signal demonstrates both the statistical robustness and biological reproducibility of the learned influence profiles. This explicit and reproducible encoding of helical phasing highlights the power of modeling regulatory syntax as an interpretable parameter rather than an emergent property.

### 2.3 Helical phasing encodes Nanog’s long-range cooperative grammar

To further quantify the periodic signal reproducibly captured in certain motif influences learned by InfluenceNet, we performed a Fourier spectrum analysis. This approach decomposes the spatial influence functions into their constituent frequency components, allowing us to identify any dominant periodic signals embedded in the learned parameters. A sharp, dominant peak emerged at 10.5 bp in Nanog models across several motifs (Fig. 3a), matching the helical pitch of B-form DNA. Consistently, the influence-reweighted motif activations revealed helical binding preferences of Nanog that were entirely absent from the raw motif activations (Fig. 3b). This feature indicates that the model captured a consistent rotational phasing of Nanog binding with respect to the DNA helix — a signature previously associated with cooperative binding events constrained by geometric alignment along the DNA backbone.

**Figure 3:**
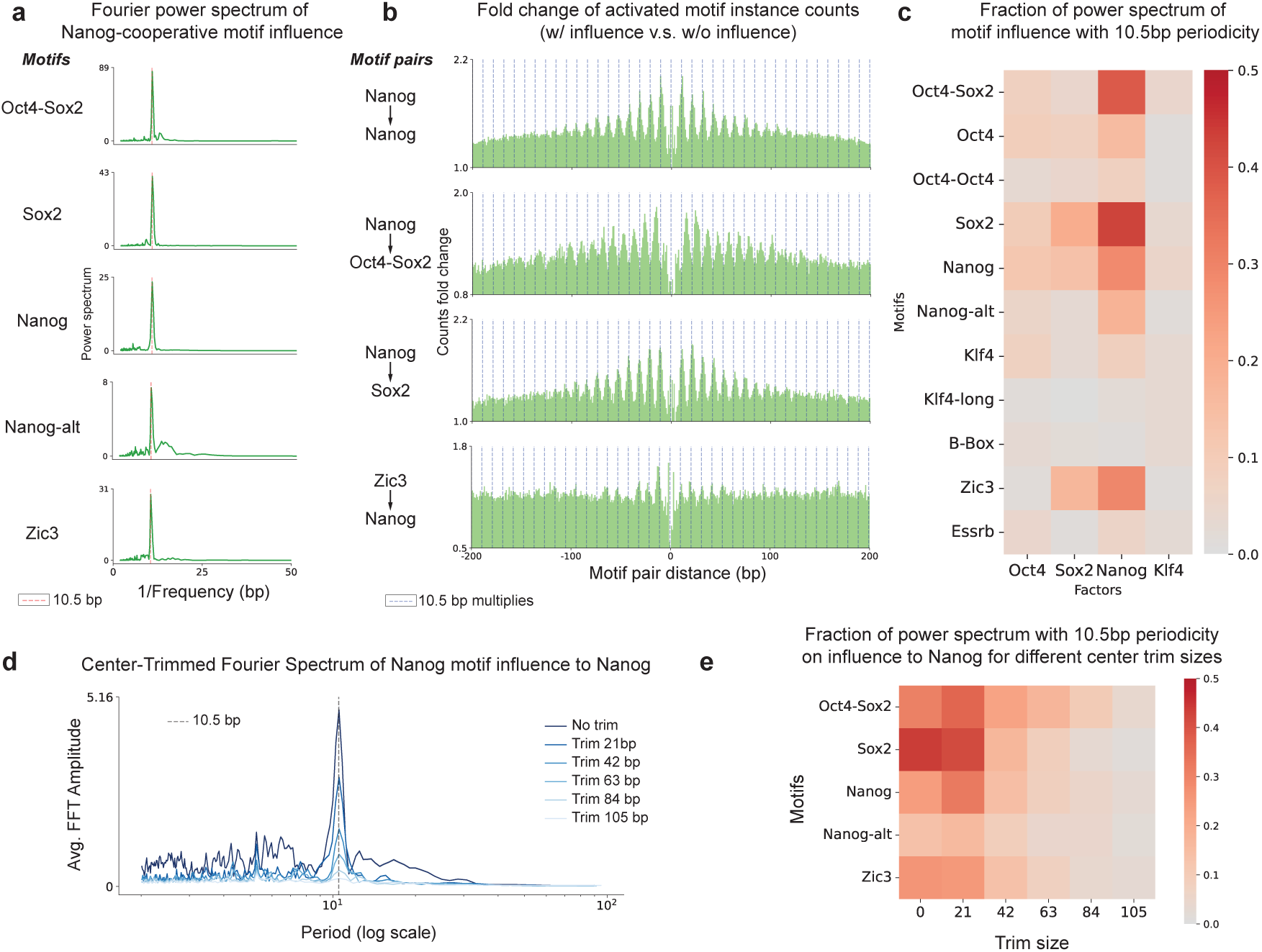
Motif influence analysis revealed 10.5-bp periodic binding of Nanog. **a**, Fourier power spectrum of averaged motif influence on Nanog binding reveals significant helical periodicity (10.5 ± 0.3bp). Low-frequency components were removed by subtracting the smoothed signal. **b**, Fold change in co-activation frequencies of motif pairs across different spacings, with or without weighting of influence on Nanog. Pronounced 10.5-bp periodicity is observed only when motif influence is applied. **c**, Fraction of power spectrum with 10.5-bp periodic component on averaged motif influence. **d**, Center-trimmed Fourier analysis of motif influence on Nanog binding shows that the 10.5-bp periodicity decreases gradually with increased trimming. **e**, Heatmap illustrating that the strength of helical periodicity diminishes at varying distance for different motifs.

When we summarized the prevalence of 10.5-bp frequencies across all motif-assay combinations, Nanog stood out as showing the strongest and most coherent helical periodicity. Sox2 displayed weaker but detectable oscillations, Oct4 only faint traces, and Klf4 none at all (Fig. 3c; all Fourier spectra in Supplementary Fig. 5b). This gradient mirrors the distinct binding behaviors of these factors: Nanog’s tendency to engage DNA cooperatively at phased sites, Sox2’s partial helical sensitivity due to its flexible HMG-domain binding mode, and Klf4’s largely independent, site-specific binding. Importantly, because the periodicity arises directly from model parameters rather than simulation-based perturbations, they represent an explicitly encoded and highly reproducible feature of the learned grammar, not a byproduct of downstream analysis.

Interestingly, the periodic influence in the Nanog assay extended to spacings approaching 100 bp, well beyond the range of direct protein-protein contacts. To quantify the spatial reach of this phasing, we performed Fourier analyses on influence profiles with progressively trimmed central regions in multiples of 10.5 bp (Fig. 3d; summarized in Supplementary Fig. 6). The 10.5-bp signal persisted up to 84 bp for Oct4-Sox2 and 63 bp for Nanog, Nanog-alt, and Zic3, suggesting that helical coherence is maintained over several turns of the DNA helix Fig. 3e).

This observation is striking because strict 10.5-bp periodicity implies a high degree of geometric alignment between DNA-bound complexes, yet the persistence of this coherence over such distances exceeds expectations for simple, rigid protein-protein interactions. One plausible explanation is that Nanog forms higher-order, DNA-mediated assemblies that preserve helical phase through cooperative oligomerization. This interpretation is consistent with biochemical evidence showing that Nanog’s WR (tryptophan repeat) domain mediates prion-like multimerization on DNA [22]. Although our modeling does not directly establish this mechanism, the reproducible and long-range periodic influences learned by InfluenceNet are fully consistent with a model in which Nanog cooperativity arises from oligomeric binding arrangements that maintain helical geometry over extended genomic distances.

### 2.4 De novo motif discovery offers additional insights into Nanog cooperativity

To investigate the sequence patterns captured by InfluenceNet, we analyzed the filters from its first convolution layer. We first assessed the 11 representative motifs that were pre-initialized before training. Their convolutional weights remained largely unchanged during training (Supplementary Fig. 8), indicating that these motifs were already strongly predictive of binding of the four TFs and required minimal optimization. Other than these preinitialized motifs, InfluenceNet learned additional de novo motifs that contribute to TF binding prediction. We performed a similar analysis as AI-TAC [23] for identifying reproducible motif patterns from the first convolution layer filters. Briefly, we extracted short sequences that strongly activated each filter, computed their position frequency matrices (PFMs), and derived PWMs. We retained only motifs with high information content and reproducibility across at least five independent models (see Methods).

We used TomTom [24, 25] to cluster similar motifs, and found 7 major clusters of learned de novo motifs. In our model, each motif is associated with two additional kernels – the influence kernel which captures contextual interactions with other sequence features, and the footprint kernel which captures assay-specific signals without intrinsic biological meanings. A scatter plot of footprint height versus influence height revealed distinct motif behaviors (Fig. 4a). In the upper left corner, motifs in Cluster 2 (poly-C/G) and part of Cluster 3 (AT-rich) displayed strong footprints but weak influence, suggesting that these patterns primarily reflect assay-specific sequence biases with no actual biological meanings.

**Figure 4:**
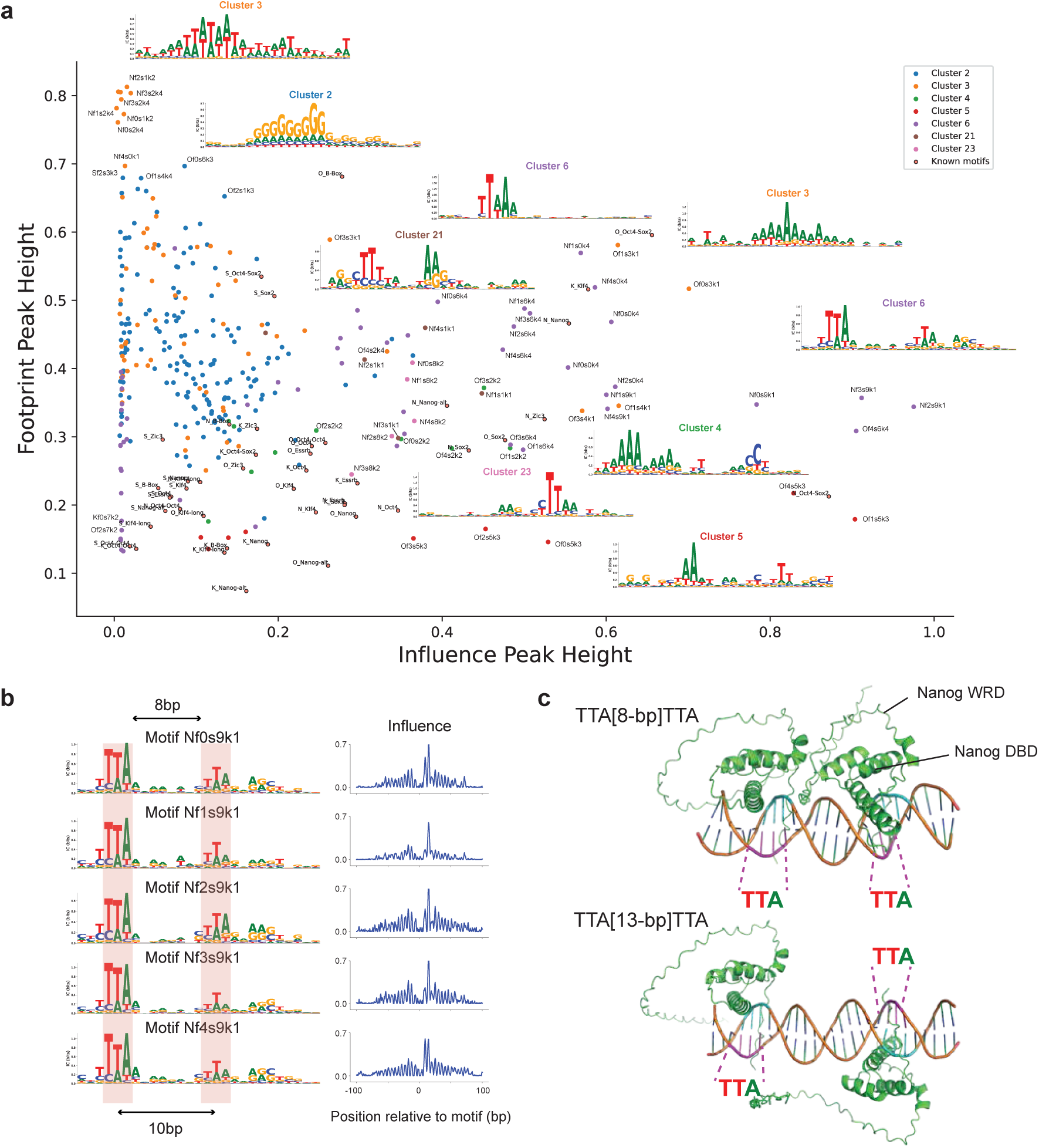
De novo motif patterns learned by InfluenceNet reveal Nanog-Nanog cooperative binding geometry. **a**, Scatter plot of footprint height versus influence height for reproducible de novo motifs (detected in ≥ 5 free kernels across 10 random seeds and 5 data splits). Motifs were grouped into 7 clusters based on PWM similarity. A representative motif logo is shown for each cluster. Node labels represent the corresponding factor (’O’-Oct4, ‘S’-Sox2, ‘N’-Nanog, ‘K’-Klf4), data split, random seed, and kernel id. **b**, Example Nanog-specific motif from Cluster 6, containing two “TTA” subsequences separated by 10 bp, or with a 8-bp spacer. This pattern shows helical periodicity in its influence profile, consistent with cooperative Nanog binding. **c**, Example AlphaFold3-predicted structures of two Nanog proteins (each comprising a DNA-binding domain and a tryptophan repeat domain) bound to “TTA…TTA” motifs. The upper structure illustrates two Nanog proteins bound on the same side of the DNA helix (8 bp spacer, i.e. one helix turn), while the lower structure shows binding on opposite sides (13 bp spacer, i.e. one and a half helix turn).

In contrast to these likely assay-derived patterns, several other motif clusters exhibited high influence with comparatively weak footprint signal, indicating biologically cooperative effects that are not the result of direct transcription factor binding. We identified two Nanog-specific clusters (Cluster 6, Cluster 23) sharing a common “TTA” subsequence, which has some overlap with previously identified Nanog motifs [26, 27]. Cluster 6 is especially notable: it is highly reproducible across independent trainings, and contains of two short “TTA” subsequences separated by 10 bp (Fig. 4b). This spacing matches the 10.5 bp helical periodicity that we observed for Nanog-associated influence profiles (Fig. 2c, 3), suggesting that Cluster 6 may represent a cooperative Nanog-Nanog binding grammar in which two weak half-sites are positioned on the same face of the DNA helix.

To evaluate whether this putative Nanog-Nanog grammar is geometrically plausible, we modeled complexes of two Nanog molecules bound to DNA using AlphaFold3 [28] (see Methods). We provided two DNA substrates: one containing a “TTA…TTA” pattern with an 8 bp spacer we observed in our model, and one with a 13 bp spacer. For each DNA substrate we modeled two copies of Nanog comprising its homeodomain and the C-terminal tryptophan repeat (WR)/oligomerization region. Across multiple predictions, AlphaFold3 consistently placed the two Nanog homeodomains on the same side of the DNA helix for the 8 bp spacer, but on opposite sides for the 13 bp spacer (Fig. 4c, more examples in Supplementary Fig. 9).

We also note that the oligomerization WR domains were as expected unstructured and have low AlphaFold3 confidence scores. However, we retained them in the model to enable downstream molecular dynamics simulations aimed at quantifying the interaction potential between Nanog molecules in different configurations.

### 2.5 Molecular modeling supports a plausible mechanism for Nanog oligomerization

InfluenceNet analysis revealed a distinct 10.5-bp helical periodicity in Nanog’s influence profiles and identified a de novo motif featuring two short “TTA” half-sites separated by approximately 10 bp. AlphaFold3 modeling predicted that two Nanog monomers bound to such a motif occupy the same face of the DNA helix at 8 bp spacing, but opposite faces at 13 bp spacing. However, AlphaFold3 cannot reliably model the flexible C-terminal WR region, which has been shown biochemically to mediate Nanog self-association [22]. To evaluate whether the predicted geometry could support direct or indirect inter-monomer interactions, we performed all-atom molecular dynamics (MD) simulations of both configurations. Each simulated system consisted of two Nanog monomers bound to a 25-bp double-stranded DNA fragment containing either an 8-bp or 13-bp TTA-TTA spacing. The protein construct encompassed both the DNA-binding domain (DBD) and the WR region, while omitting the remainder of the N-terminus for tractability.

Trajectory analysis confirmed that the overall DNA conformation and relative DBD positioning remained stable throughout the simulations, whereas the WR regions displayed high mobility, consistent with their intrinsic disorder. As expected, contact-map analysis revealed substantially closer inter-monomer residue distance in the 8-bp spacer configuration, indicating that the shorter spacer favors transient encounters between the Nanog monomers (Fig. 5b). Unexpectedly, the most recurrent and stable contacts occurred between the WR region of one monomer and the DBD of the other, rather than symmetric WR-WR pairing (Fig. 5b, c). The representative configuration in Fig. 5a shows that the second monomer’s WR tail remains solvent-exposed, potentially available to engage an additional monomer.

**Figure 5:**
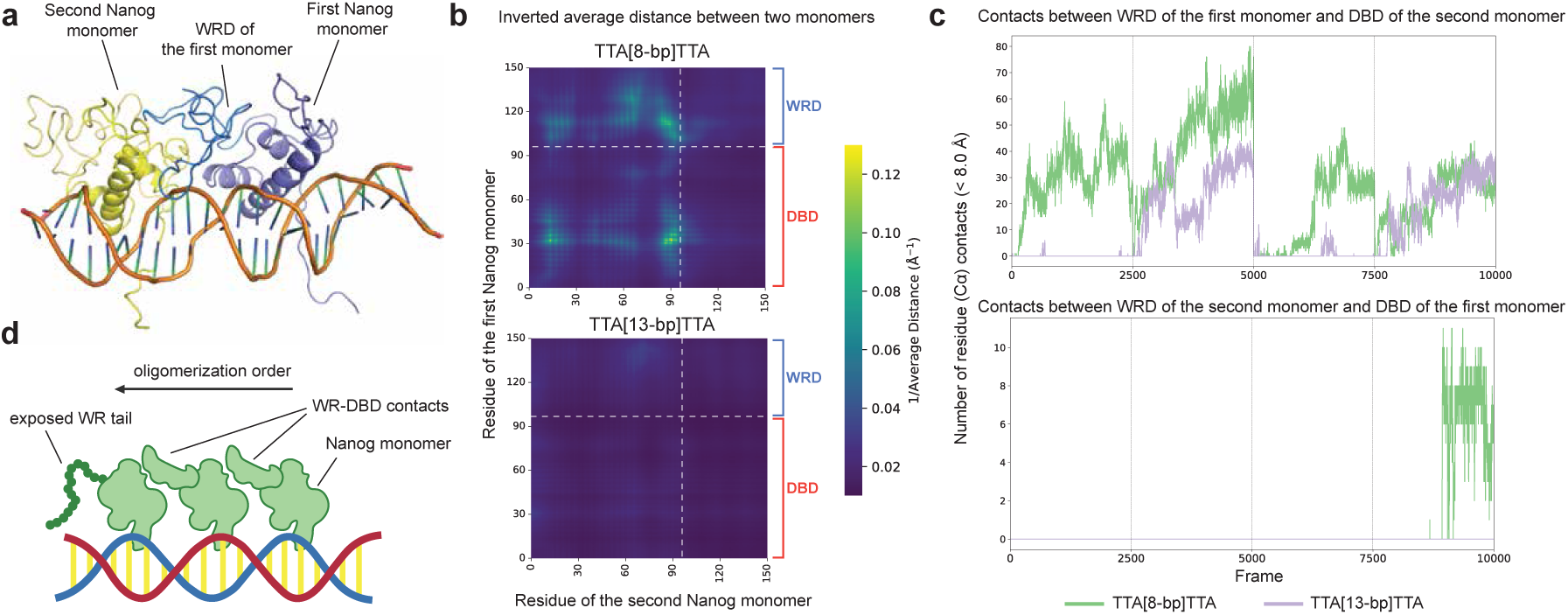
MD simulations supports chained oligemerization of Nanog monomers. **a**, A representative MD snapshot of the two Nanog monomers dimerizing asymmetrically through WRD-DBD contacts. **b**, Inverted average residue distance map showing closer monomer-monomer proximity in the 8-bp spacer system relative to 13 bp. **c**, WRD-DBD number of contacts (< 8.0 Å) between the first and second Nanog monomers for both directions. The 8-bp spacing significantly enhances contact frequency. Second monomer contacts are less frequent, supporting an assymetric configuration. **d**, Schematic model of chain-like oligemerization, in which of one monomer interacts with the DBD of the next, maintaining helical alignment along the DNA.

These findings suggest that Nanog could assemble on DNA through sequential, asymmetric WR-DBD contacts, forming chain-like oligomers rather than symmetric dimers (Fig. 5d). Such chaining naturally preserves helical phase alignment and could propagate over tens of base pairs, consistent with the long-range, periodic cooperativity inferred from the learned influence kernels from InfluenceNet.

Although these simulations do not constitute experimental evidence for a stable multimer, they provide a physically plausible model consistent with (1) the learned 10.5-bp periodic influence pattern, (2) the static AlphaFold3 geometry, and (3) prior biochemical evidence of WR-mediated self-association. Together, these results suggest a structural hypothesis in which Nanog’s disordered WR domains mediate weak, asymmetric bridging between adjacent monomers, forming higher-order DNA-bound assemblies that maintain helical register and may underlie Nanog’s spacing-sensitive cooperativity.

## 3 Discussion

InfluenceNet provides a transparent framework for modeling transcription-factor binding by directly parameterizing motif influence, enabling mechanistic interpretation without post-hoc perturbations. Despite its simplicity, the model achieves predictive accuracy approaching that of high-capacity dilated CNNs, demonstrating that much of TF binding behavior can be captured through interpretable, factorized components. At the same time, the modest performance gap highlights that standard deep architectures may extract additional nonlinear dependencies that are not readily represented in the current formulation. It remains unclear whether this additional learned complexity reflects meaningful biological effects or simply captures assay-specific sequence biases, as suggested by the structured yet assay-dependent patterns observed in the footprint kernels.

The explicit representation of motif influence also revealed both expected and novel forms of cooperativity. The sharp, reproducible 10.5-bp periodicity observed for Nanog influence profiles provides compelling evidence that the model captures the helical geometry of cooperative binding. The much weaker periodic signatures in Oct4 and Sox2, and their absence in Klf4, are intriguing and may indicate subtle or transient modes of phase-sensitive cooperativity that warrant further investigation.

Finally, our structural modeling and molecular-dynamics simulations propose a mechanistic explanation for Nanog’s long-range cooperativity — sequential, asymmetric WRD-DBD contacts that allow chain-like oligomerization along DNA. While this model aligns with both our learned influence profiles and prior biochemical observations, direct experimental validation such as cross-linking or single-molecule assays are required to confirm this mode of assembly.

Together, these findings illustrate how explicitly interpretable architectures like InfluenceNet can bridge predictive modeling and mechanistic hypothesis generation, offering a scalable framework for dissecting the syntax of cis-regulatory grammar. Our implementation of InfluenceNet and corresponding analysis scripts are available at https://github.com/chikinalab/InfluenceNet.

## 4 Methods

### 4.1 ChIP-nexus data and preprocessing

The experimental strand-specific ChIP-nexus data used in this study is from BPNet [3] for four pluripotent TFs: Oct4, Sox2, Nanog and Klf4 in mouse embryonic stem cells. Signal tracks were smoothed using a 10 bp moving-average window to reduce noise. We applied an arcsinh transformation to the signal tracks to further stabilize variance. Reverse-complement sequences and binding profiles were generated as augmented training examples to account for reverse-complement equivariance of ChIP-nexus readouts.

### 4.2 InfluenceNet architecture

InfluenceNet takes local DNA sequence as input and predicts strand-specific base-pair resolution ChIP-nexus profile. The surrounding ±500-bp DNA sequence is one-hot encoded into a 1000 × 4 matrix. The main body of InfluenceNet consists of three convolution-based modules: the sequence, influence, and assay layers. The sequence layer scans the 1000-bp input sequence to detect motif patterns. It uses 26 filters of width 27. Among them 22 are initialized with position-specific scoring matrices (PSSMs) from 11 representative motifs identified by BPNet [3] and their reverse complements, while 4 are randomly initialized to allow de novo motif discovery. This layer outputs raw motif activations based purely on local sequences. The influence layer models long-range cooperative effects between motifs using 26 filters of width 401, implemented via FFT-based convolution for computational efficiency. The influence layer is followed by a fully-connected linear layer (26 × 26) that aggregates all influences in weighted sums. Transformed motif influences are multiplicatively applied to the raw motif activations to yield influenced motif activations. Finally, the assay layer maps these activations to ChIP-nexus profiles through filters of width 201, together with a final linear transformation (26 × 4). For strand-specific prediction, two independent assay and output layers are used for two strands, while the upstream sequence and influence layers are shared.

### 4.3 InfluenceNet training

InfluenceNet was trained to predict ChIP-nexus profile for one or multiple TFs depending on the experimental settings. For performance comparison in the 5-fold cross-validation and held-out test, we trained two separate models to predict forward and reverse strand profiles for all four TFs, consistent with the BPNet configuration. To examine biologically interpretable parameters learned by InfluenceNet, independent models were trained for each TF, such that learned parameters for different TFs are not mixed up. Models were optimized using mean-squared error (MSE) between predicted and observed read counts at each nucleotide position. To regularize the influence layer, we applied L1 weight regularization and L2 smoothness constraints between spatially adjacent kernel weights. Models were implemented using pytorch v.2.6.0 and trained with the Adam optimizer (learning rate = 0.001). Early stopping for training was triggered after 3 epochs without improvement in validation performance. Models were trained with a batch size of 256.

### 4.4 Comparison with BPNet

For benchmarking, we reimplemented BPNet using pytorch, replicating its architecture and hyperparameters as described in the original publication [3]. The BPNet architecture we used consisted of the standard convolutional body (with 9 dilated convolution layers) followed by a single 1D convolutional output head with 4 channels corresponding to the 4 TFs. InfluenceNet and BPNet were trained under identical data splits, loss functions, and optimization schemes to ensure a fair comparison. Model performance was evaluated using two metrics: 1) the Spearman correlation between predicted and observed profiles, for evaluating the accuracy of profile shape prediction, and 2) the cross-region correlation, defined as the Spearman correlation between predicted and observed total read counts across sequence regions, used for evaluating the accuracy in total signal counts prediction. The two metrics were averaged across data folds and strands to obtain the final reported performance.

### 4.5 Identification of reproducible motif patterns learned by the first convolution layer

We followed a similar approach as AI-TAC [23] to identify sequence motifs captured by the first convolutional layer of InfluenceNet. First, samples with reliable predictions (correlation > 0.2 between outputs and targets) were selected, and we extracted activations from the free kernels. All 27-bp sequence regions activated by at least 50% of the maximum activation per filter were aligned to construct PFMs, which were then converted into PWMs. We preserved only the reproducible motif PWMs that were captured by at least five independent models, and with high information content (> 0.4). To obtain similarity-based clustering on the identified motifs, we used Tomtom [24, 25] to compute similarities and p-values. A graph-based clustering approach was then applied, where motifs were connected if their pairwise similarity was significant (with p-value < 0). Connected components within this network were identified as motif clusters.

### 4.6 Predicting Nanog binding to de novo motifs using AlphaFold3

To examine the binding configurations of Nanog to the de novo motif pattern characterized by 10 bp-spaced TTA subsequences, we used Alphafold3 to predict the structures of two Nanog molecules bound to the corresponding DNA consensus sequence. Specifically, we generated multiple random consensus DNA sequences containing a a “TTA…TTA” pattern. The amino acid sequence of Nanog obtained from Uniprot [29] (Q9H9S0), comprising its homeodomain and the C-terminal WR region (residue 91-240), was used as input, with two copies included for examining their relative orientation to the DNA sequence. Each consensus sequence and its reverse complement were also provided to AlphaFold3. The predicted structures were visualized using PyMol (v3.0.0).

### 4.7 Molecular dynamic simulations of the Nanog-Nanog dimer

The protein-DNA complex was simulated using AMBER 24 [30]. The system was neutralized with sodium ions and solvated using OPC water in an octohedral box with 12Å padding. The OPC water model [31] was selected due to its reported improved performance when simulating intrinsically disordered proteins [32] such as the C-terminal of Nanog. The ff19SB protein force field [33] was used with the Parmbsc1 [34] nucleic acid force field. Initial equilibration consisted of two minimization steps (the first with protein restraints), followed by 1 ns of restrained NVT dynamics (heating from 0 K to 300 K), and 1 ns of unrestrained NPT dynamics at 300 K and 1 atm. Four 250ns production runs (NPT at 300K and 1 atm) were performed for each system. Analysis was performed using MDAnalysis [35].

## 5 Supplementary Information

### 5.1 Supplementary Figures

**Supplementary Figure 1:**
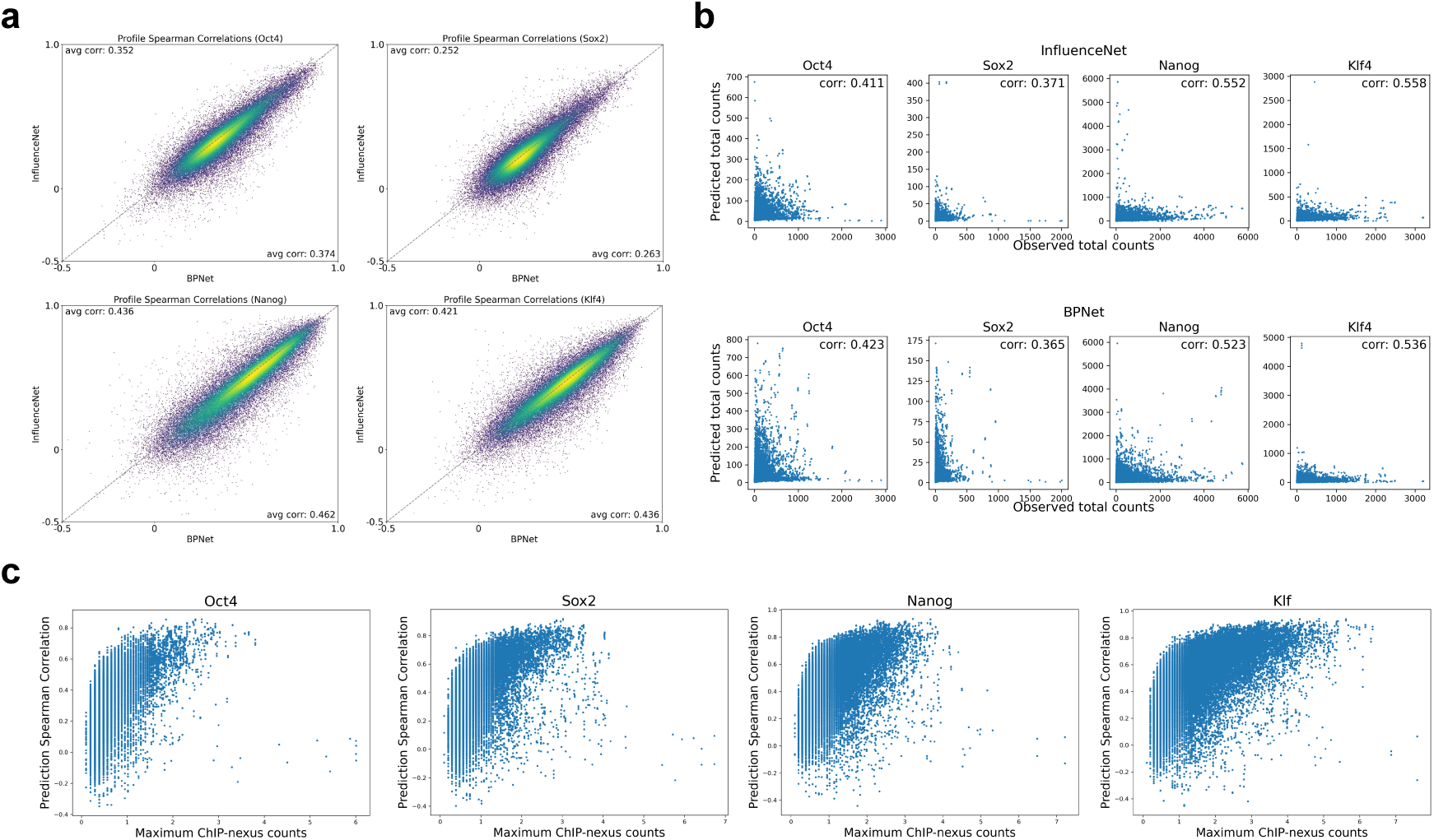
Comparison of InfluenceNet with BPNet performance on held-out test chromosomes. **a**, Scatter plot comparing BPNet (x axis) and InfluenceNet (y axis) prediction profile Spearman correlations on the held-out test chromosomes. **b**, Comparison of total counts prediction correlations for BPNet (top row) and InfluenceNet (bottom row). **c**, The maximum ChIP-nexus counts per profile in the test dataset on the x axis vs. the InfluenceNet profile prediction Spearman correlations. The performance is highly correlated with the profile total signals. All evaluations were performed on held-out test chromosomes unseen during model training and validation.

**Supplementary Figure 2:**
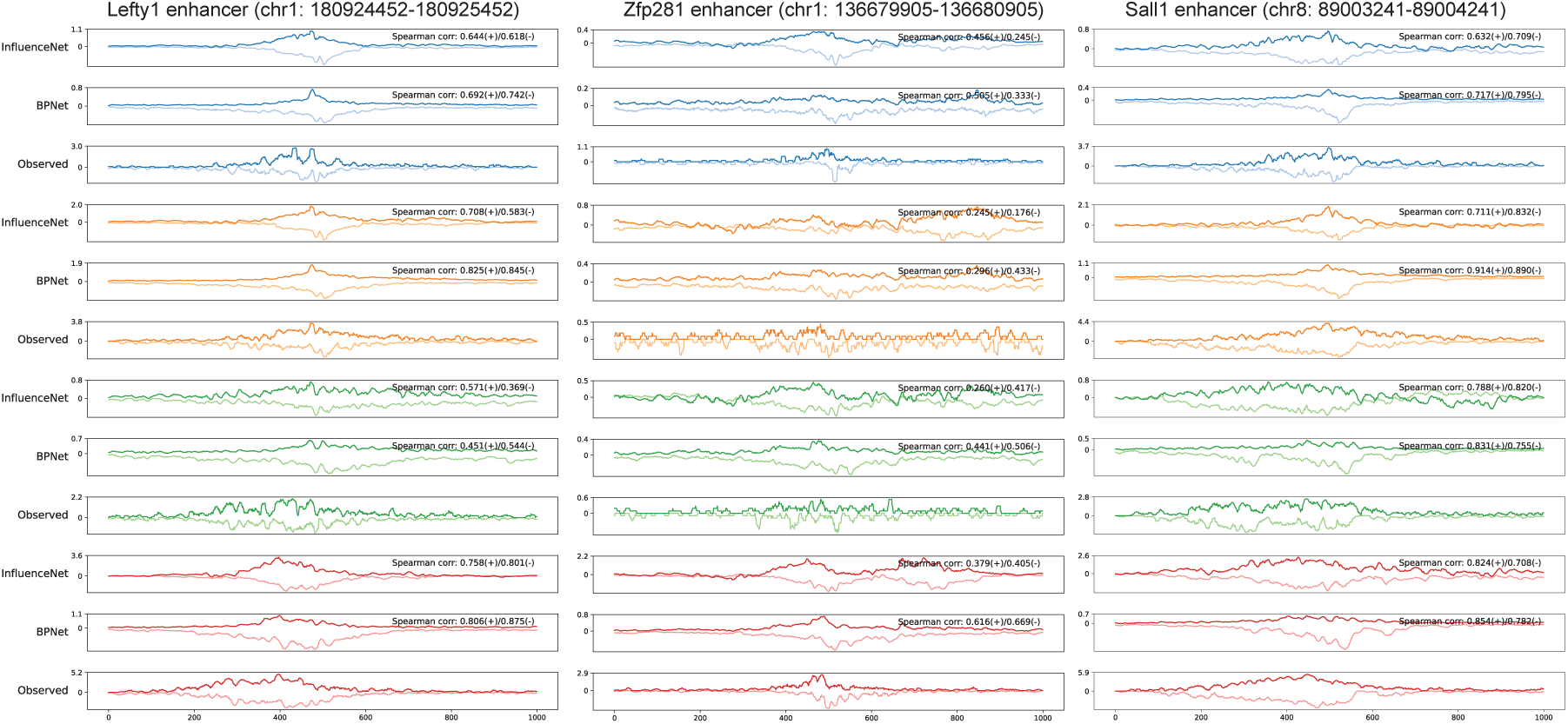
Example predictions of InfluenceNet compared with BPNet. Predictions of InfluenceNet and BPNet on example regions on held-out test chromosomes, compared with the corresponding experimental ChIP-nexus profiles. Left, Lefty1 enhancer region (chr1:180,924,452-180,925,452); Middle, Zfp281 enhancer region (chr1: 136,679,905-136,680,905); Right, Sall1 enhancer region (chr8: 89003241-89004241).

**Supplementary Figure 3:**
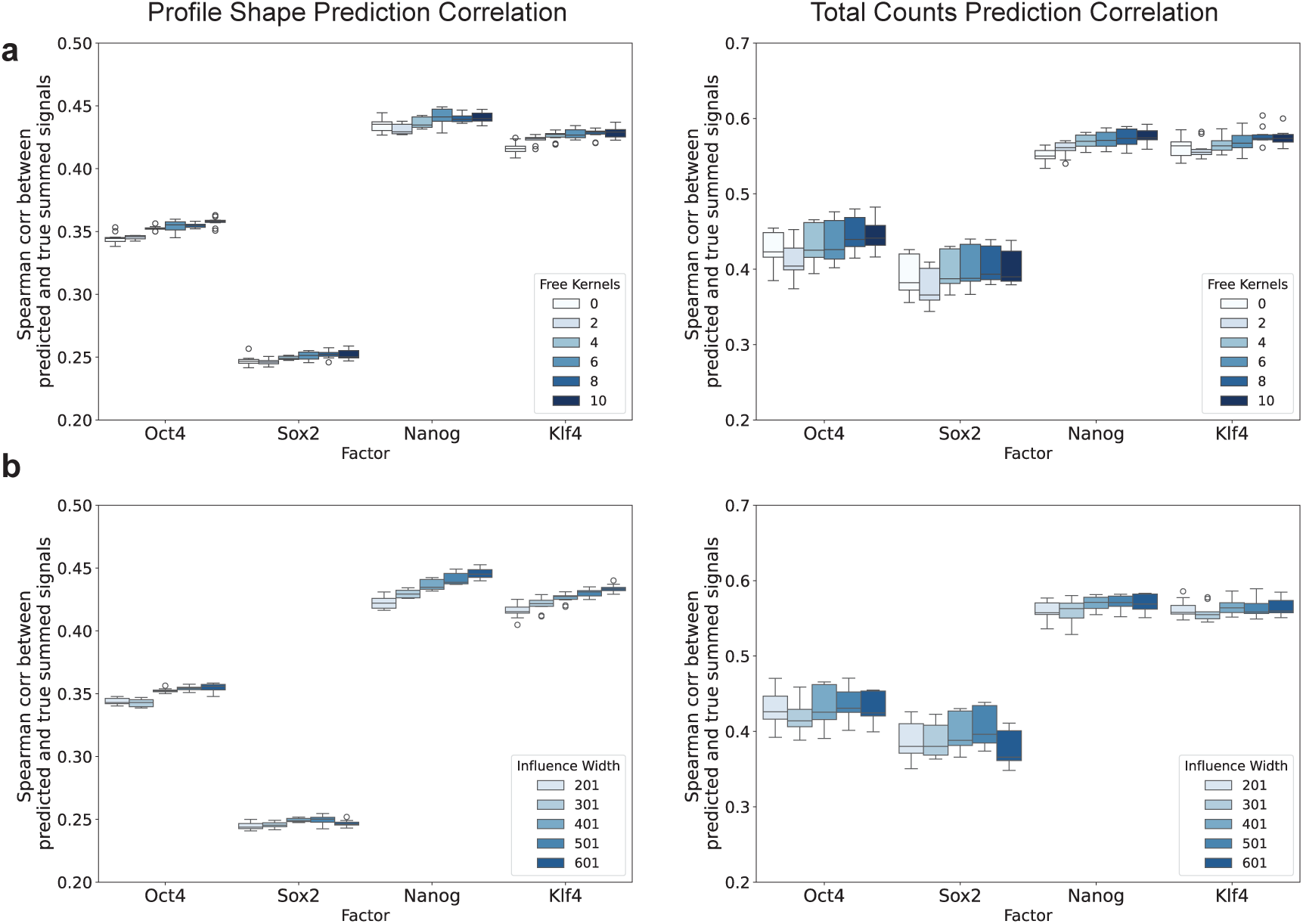
Performance of InfluenceNet across different number of free kernels and influence kernel widths. **a**, Profile shape and total counts performance of InfluenceNet with varying number of free kernels on cross-validation data. **b**, Profile shape and total counts performance of InfluenceNet with varying influence kernel widths on cross-validation data.

**Supplementary Figure 4:**
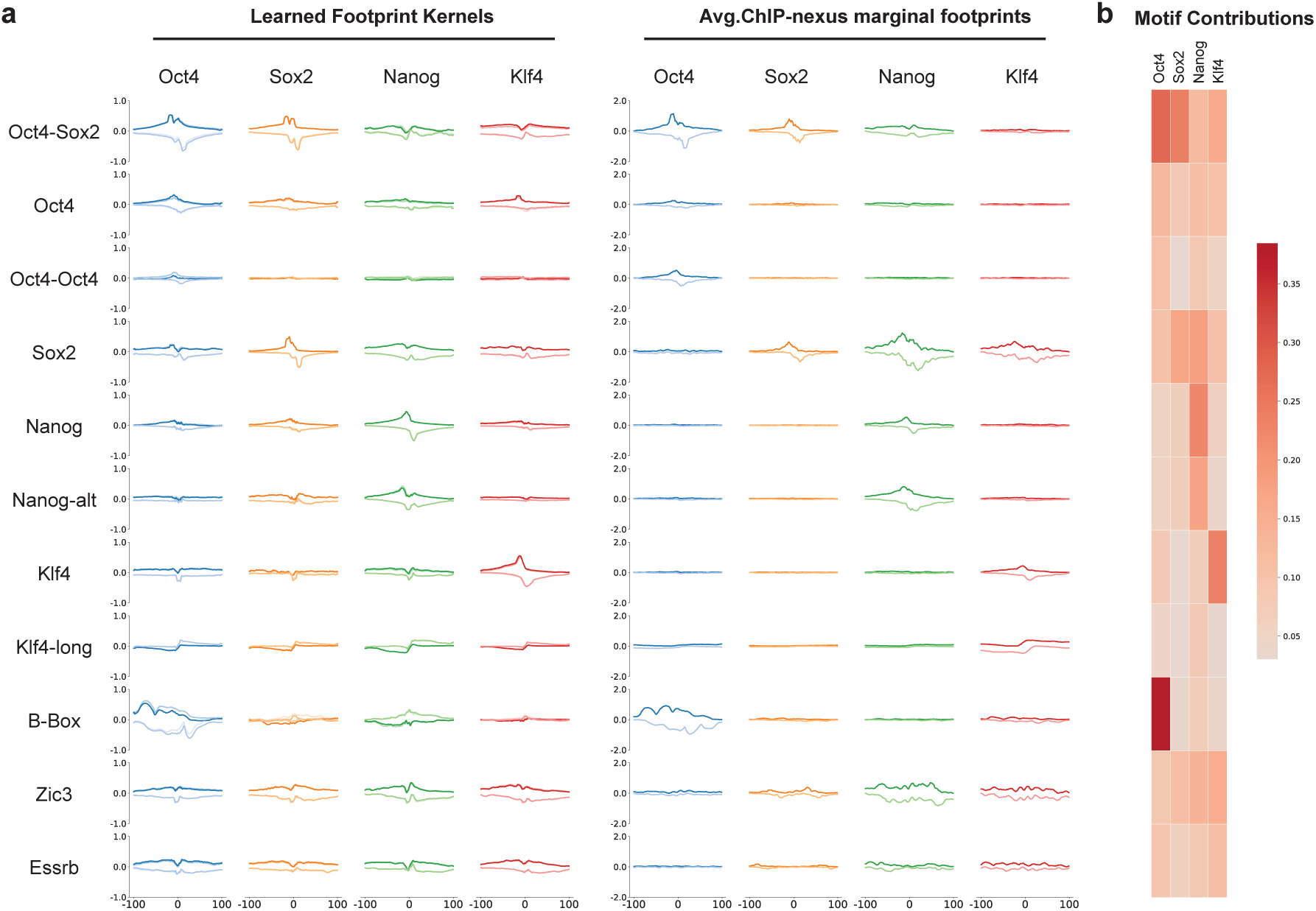
Assay layer kernels and final linear layer weights learned by InfluenceNet. **a**, InfluenceNet learned footprint kernels largely resemble experimental ChIP-nexus footprints. Left, Averaged ChIP-nexus footprints centered on the top 200 motif occurrences identified by FIMO scanning [36], ranked by their log-odds scores. Dark and light colors indicate footprints on forward– and reverse-strand motifs, respectively. **b**, Linear weights extracted from the final linear layer of InfluenceNet, which computes a weighted sum of motif-specific footprints to generate the eventual binding profile. The heatmap highlights motifs with strong contributions to each TF’s predicted binding signal.

**Supplementary Figure 5:**
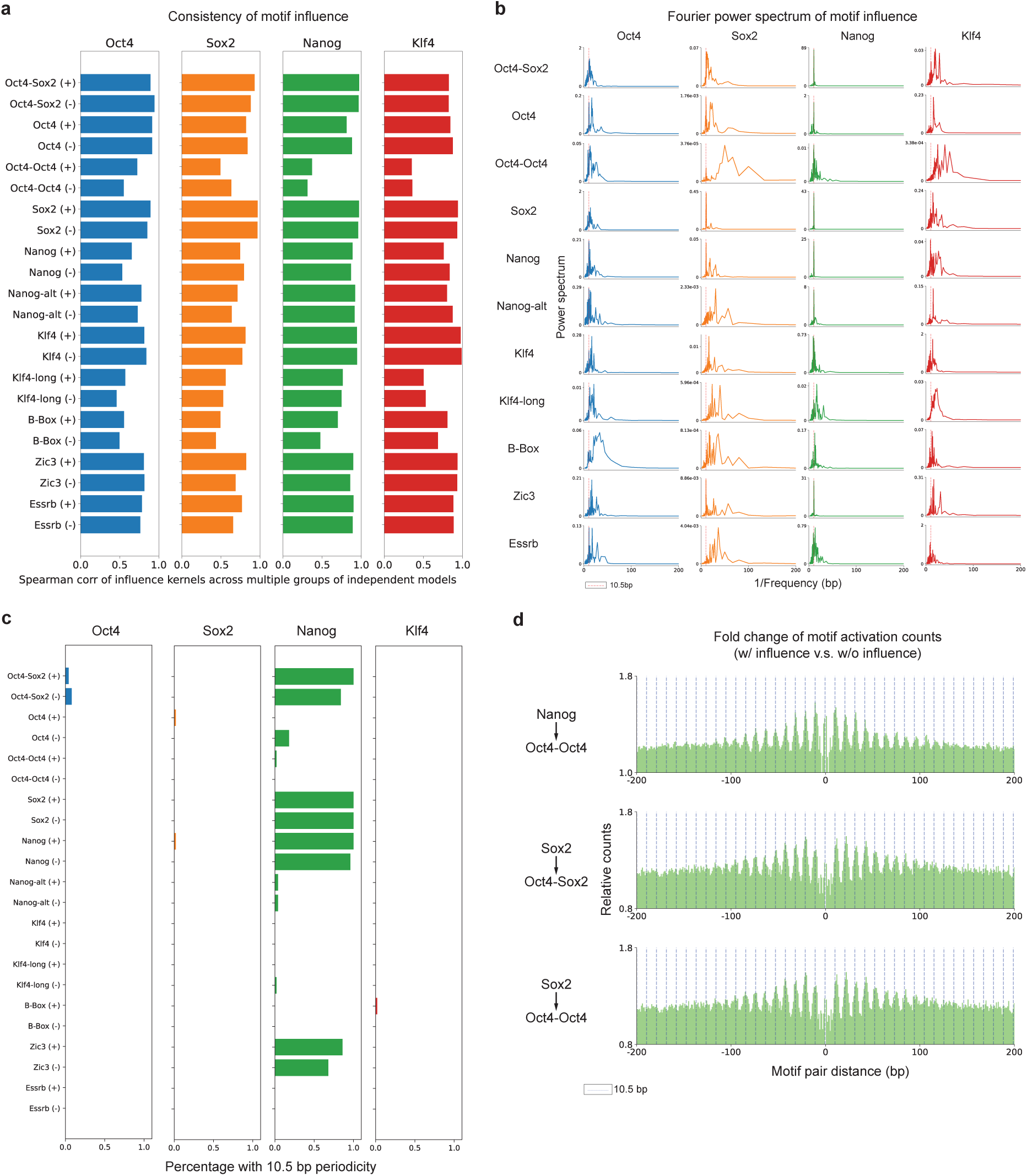
Periodicity and consistency of motif influence kernels. **a**, Strongly influential motifs showed highly consistent influence profiles across independent model replicates, whereas weakly influential motifs displayed variable or noisy patterns, as measured by replicate Spearman correlations. **b**, Fourier power spectra of averaged motif influence. A distinct 10.5-bp periodicity was observed exclusively for influence on Nanog binding. Low-frequency components were removed by subtracting the smoothed signal. **c**, Frequency of motifs exhibiting a 10.5 ± 0.3 bp periodic influence across 50 independent models. Nanog cooperative motifs consistently showed robust helical periodicity, whereas other motifs and factors did not. **d**, Fold change in co-activation frequencies of motif pairs across different spacings, with or without weighting of influence on Nanog. A 10.5-bp periodic signal was observed between Nanog and Oct4-Oct4, indicating a weak Nanog cooperative interaction. Motif pairs not involving Nanog also showed helical periodic spacings (Sox2→Oct4-Sox2, Sox2→Oct4-Oct4), likely due to the indirect combinatorial effect of Nanog helical phasing and its cooperative binding preferences with Oct4 and Sox2.

**Supplementary Figure 6:**
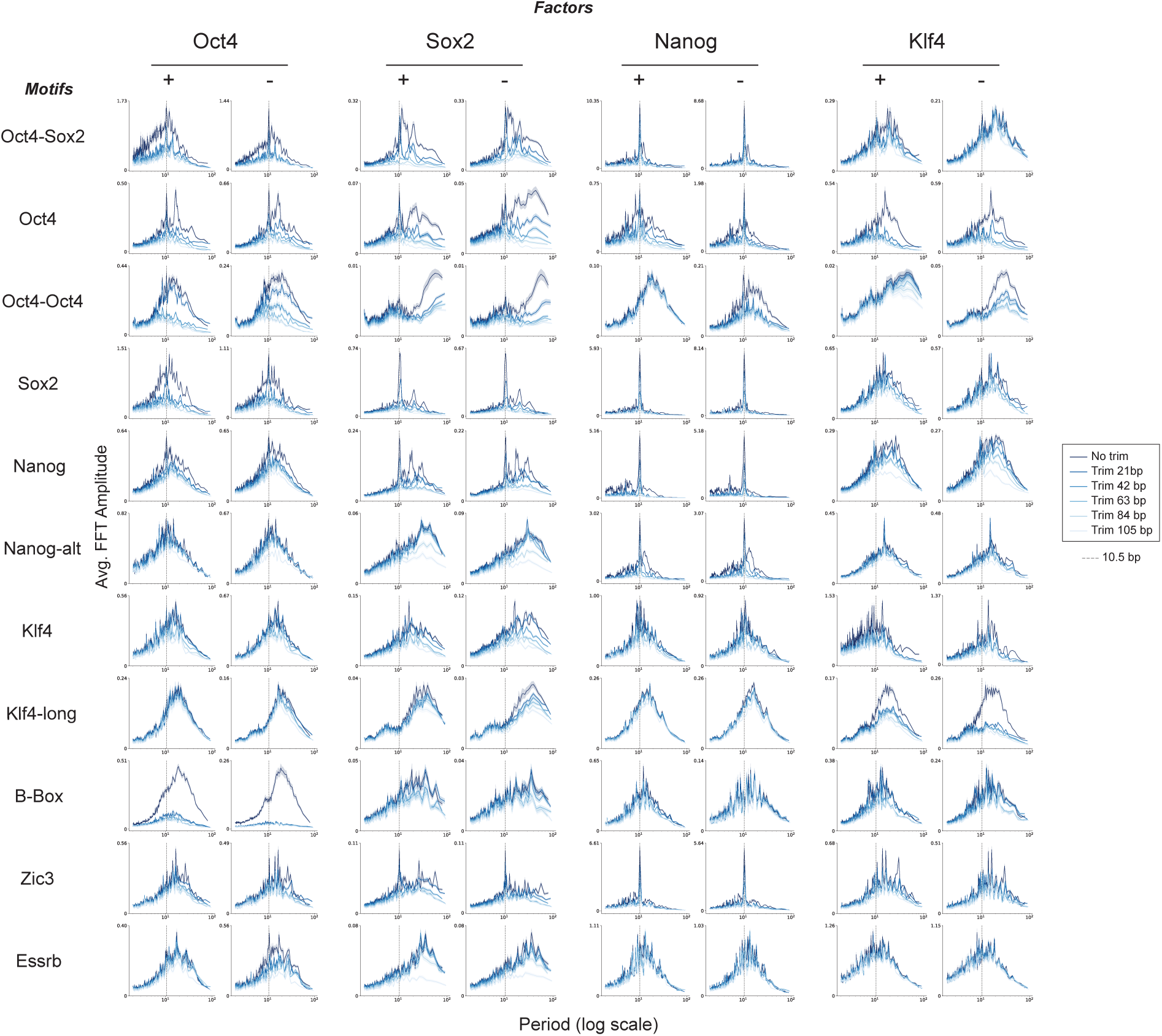
Fourier power spectra of center-trimmed motif influences. The 10.5-bp periodicity of Nanog influence persists across distances up to ±100 bp but gradually diminishes with increasing distance. For the other motifs that do not exhibit significant periodic influence, the Fourier power showed little variation across different trim sizes.

**Supplementary Figure 7:**
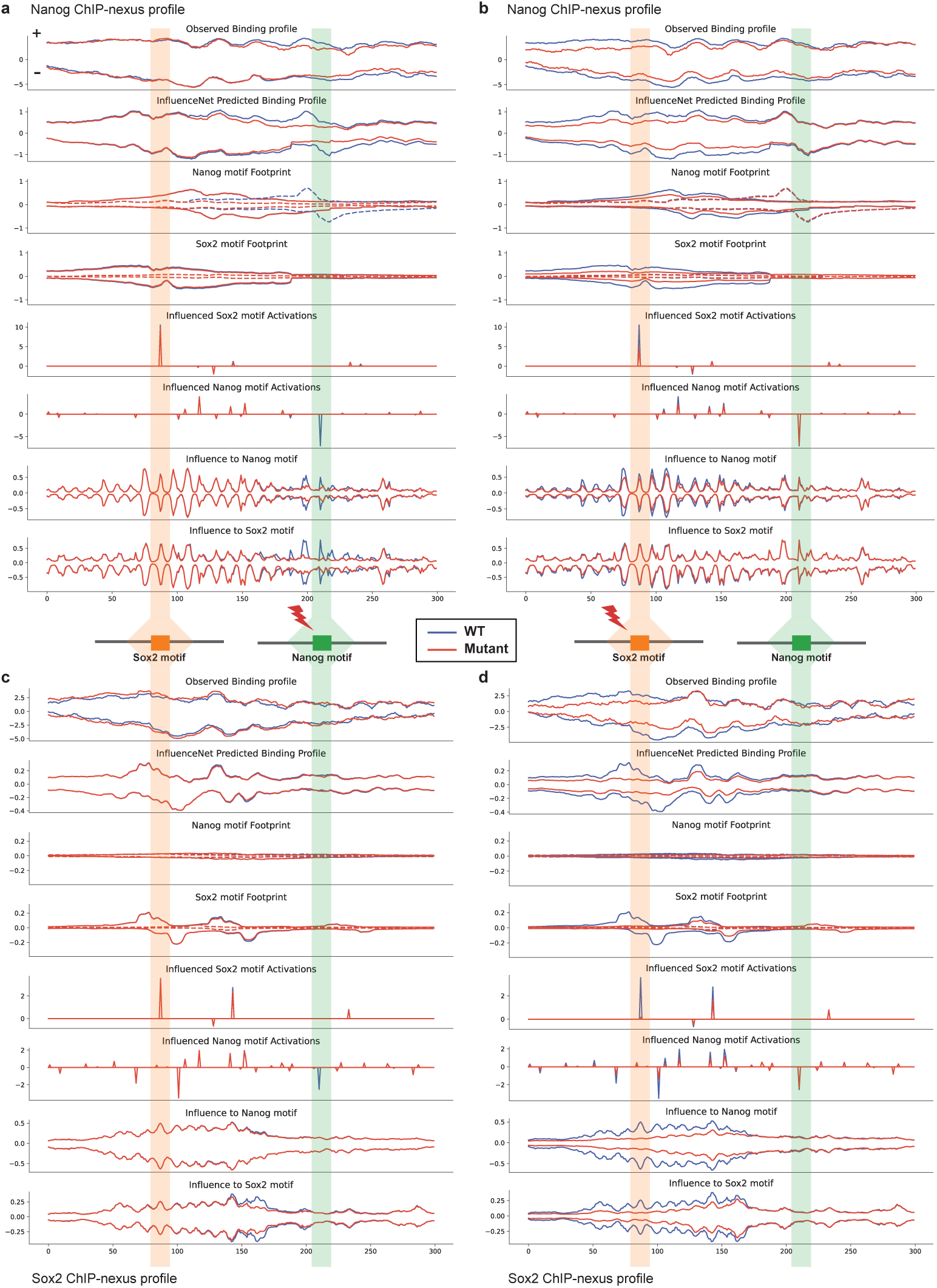
InfluenceNet predicts and explains the regulatory effects of Sox2 and Nanog motif mutations. **a-d**, InfluenceNet predictions and experimental ChIP-nexus profiles for CRISPR-induced mutations in Sox2 and Nanog motifs within a 300 bp region (chr10:85,539,550-85,539,850; data from [3]). For each case, the observed ChIP-nexus profiles are compared with InfluenceNet predictions using wild-type (blue) and mutant (red) sequences. Subsequent tracks show intermediate model outputs using wild-type and mutant inputs respectively, including motif-specific footprints (scaled by the final linear-layer weights), influence reweighted motif activation scores, and influence. Solid and dashed lines indicate forward– and reverse-strand motifs, respectively. InfluenceNet accurately predicted mutation-induced changes in binding profiles and provided mechanistic insights. Nanog and Sox2 each showed local binding loss at their respective mutated motifs (a, d). Sox2 mutation caused a pronounced reduction in Nanog binding nearby (b), whereas Nanog mutation had minimal effect on Sox2 binding (c), revealing an asymmetric Nanog-Sox2 interaction consistent with the directional influence inferred from motif influence profiles.

**Supplementary Figure 8:**
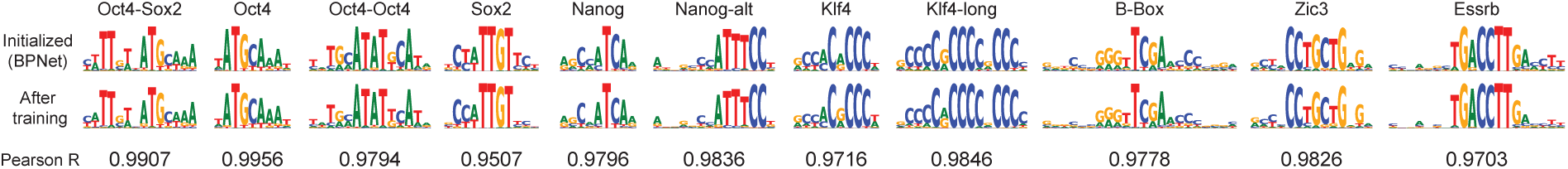
Comparison between PWMs of the 11 representative motifs before and after InfluenceNet training. Computed Pearson correlations show high robustness of the representative motif PWMs during InfluenceNet training.

**Supplementary Figure 9:**
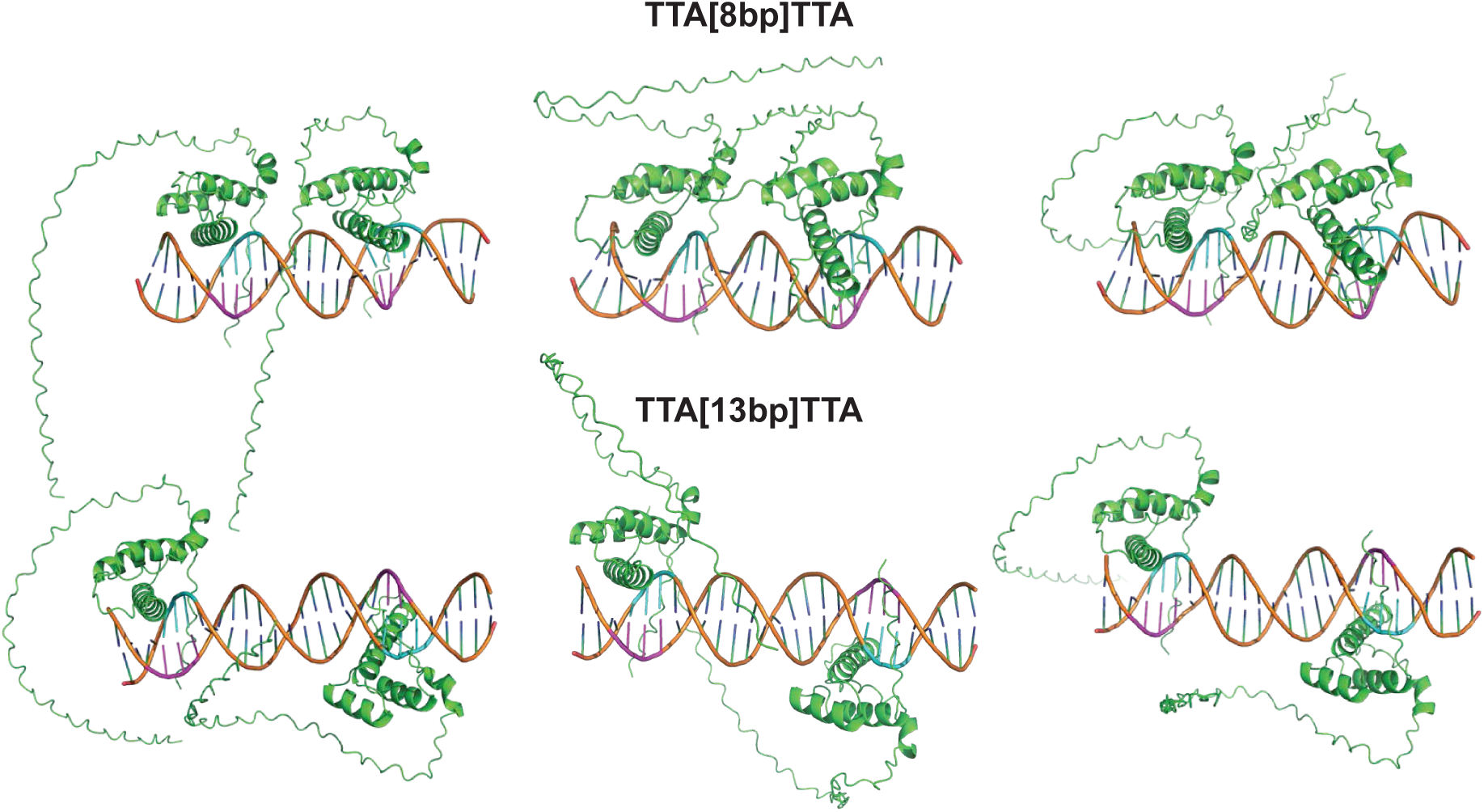
Full examples of AlphaFold3 predictions of two Nanog proteins binding to “TTA…TTA” motifs with an 8-bp spacer (first row) or with an 13-bp spacer (second row). Forward strand “TTA” subsequences are colored magneta and reverse strand “TAA” subsequences are colored cyan.

### 5.2 Supplementary Data

**Supplementary Data 1.** Reproducible clusters of de novo motifs identified by InfluenceNet. Each row in the table represents one motif instance identified through one filter of the first convolutional layer. The table has 5 columns: 1) ‘ID’ – the unique id of each motif instance, including the corresponding factor (’O’-Oct4, ‘S’-Sox2, ‘N’-Nanog, ‘K’-Klf4), data split, random seed, and kernel number; 2) ‘Seq PWM’ – motif PWM computed from the PFM aggregated over activated sequences by the filter; 3) ‘Filter PWM’ – motif PWM directly derived from the filter weight (representing learned motif PSSM); 4) ‘Influence Profile’ – the corresponding learned influence kernel; 5) ‘Footprint Profile’ – the corresponding learned footprint kernel, with dark and light colors representing footprint on the forward and reverse strand respectively.

### 5.3 Supplementary Method

#### 5.3.1 Computing averaged ChIP-nexus footprints

To compute the marginal ChIP-nexus footprints at sites of each motif, we first scanned the motif instances using FIMO [36]. We used the same PWMs from BPNet that we used for initializing kernel weights of the first convolution layer. We ran FIMO v5.5.7 on all the ±500 bp regions of ChIP-nexus peaks (same as the model inputs) using the default parameters. The output hits were ranked by the q-value from FIMO. We used the top 200 motif instances as the center positions and averaged the ChIP-nexus profiles within ±100 bp.

#### 5.3.2 Configurations of Molecular Dynamics Simulations on Nanog-DNA complexes

The AMBER molecular dynamics simulations were performed using AMBER 24 [30]. The protein-DNA complex was parameterized using the ff19SB protein force field [33] and the Parmbsc1 nucleic acid force field [34]. The complex was solvated in an octahedral box of OPC water molecules [31] with a 12 Åbuffer in all directions, and the system was neutralized by adding sodium counterions. All model building and solvation procedures were carried out with the tleap module of AmberTools24.

Prior to equilibration, the system underwent a two-stage energy minimization. In the first stage, solvent and ions were relaxed while restraining all solute heavy atoms with a 500 kcal/mol harmonic potential, allowing local optimization of the water network and ion positions. The second minimization stage released all restraints. Both minimizations used a 10 Ånonbonded cutoff under constant volume (NVT) conditions, with the steepest descent algorithm followed by conjugate gradient minimization until convergence.

Following minimization, a multi-step equilibration was performed to gradually heat and stabilize the system. The system was first heated from 0 to 300 K under constant volume (NVT) conditions using a Langevin thermostat (collision frequency γ = 1 ps^−1^). During this phase, the solute was restrained with a 10 kcal/mol potential to prevent structural distortion while the solvent adjusted around the complex. Subsequently, an isothermal-isobaric (NPT) equilibration was carried out at 300 K and 1 atm using a Berendsen barostat to equilibrate the system’s density. All equilibration steps employed a 2 fs integration timestep, with SHAKE constraints applied to all bonds involving hydrogen atoms.

The production simulations were conducted in the NPT ensemble at 300 K and 1 atm for 250 ns per trajectory, with four independent runs per system, resulting in a total of 1 µs of sampling. Temperature control was maintained using a Langevin thermostat (γ = 1 ps^−1^), and pressure was controlled by a Berendsen barostat. Nonbonded interactions were truncated at 10 Å, and long-range electrostatics were treated with the particle mesh Ewald method. Coordinates were saved every 10 ps, and restart files were written periodically to ensure recovery capability.

All trajectories were postprocessed using cpptraj and MDAnalysis [35]. The initial equilibration phase was excluded from analysis. The trajectories were centered, imaged, and stripped of solvent molecules for analysis of protein and DNA motions, hydrogen bonding, and structural fluctuations.

