## Supplemental Data 1 for "InfluenceNet: Encoding Motif Influence for Interpretable Modeling of Cis-Regulatory Syntax"

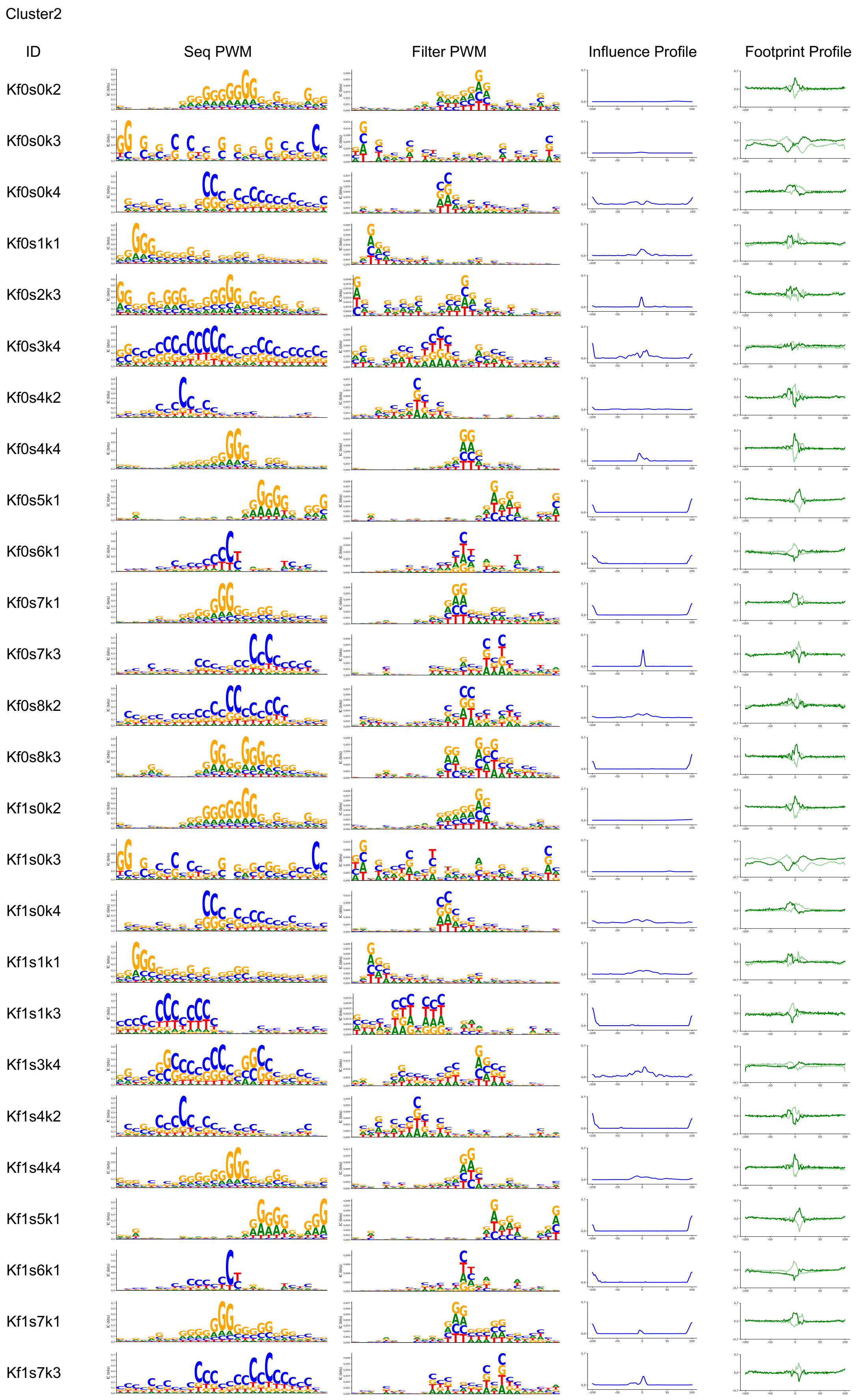
Cluster2
ID
Seq PWM
Filter PWM
Influence Profile
Footprint Profile
Kf0s0k2
Kf0s0k3
Kf0s0k4
Kf0s1k1
Kf0s2k3
Kf0s3k4
Kf0s4k2
Kf0s4k4
Kf0s5k1
Kf0s6k1
Kf0s7k1
Kf0s7k3
Kf0s8k2
Kf0s8k3
Kf1s0k2
Kf1s0k3
Kf1s0k4
Kf1s1k1
Kf1s1k3
Kf1s3k4
Kf1s4k2
Kf1s4k4
Kf1s5k1
Kf1s6k1
Kf1s7k1
Kf1s7k3

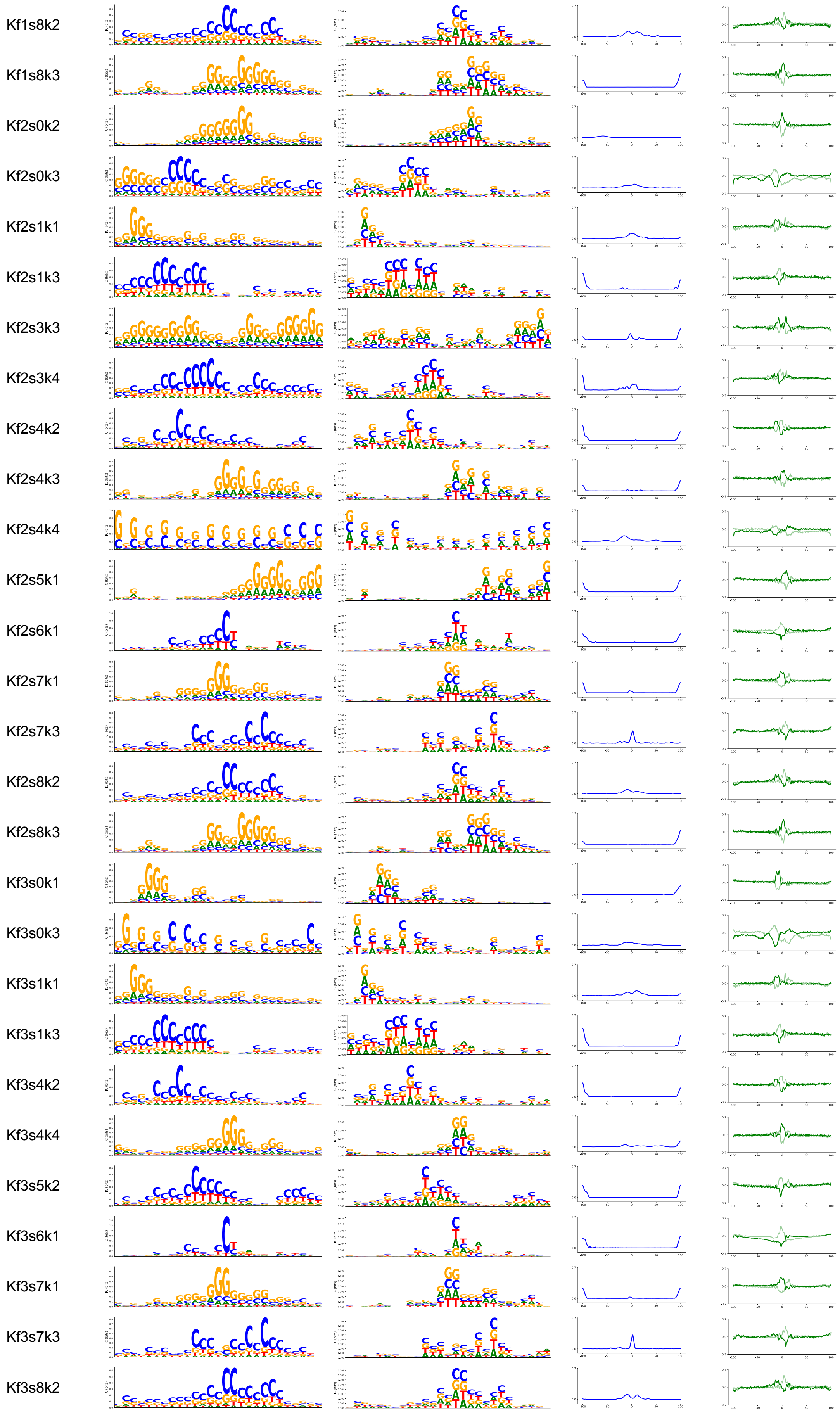
Kf1s8k2
Kf1s8k3
Kf2s0k2
Kf2s0k3
Kf2s1k1
Kf2s1k3
Kf2s3k3
Kf2s3k4
Kf2s4k2
Kf2s4k3
Kf2s4k4
Kf2s5k1
Kf2s6k1
Kf2s7k1
Kf2s7k3
Kf2s8k2
Kf2s8k3
Kf3s0k1
Kf3s0k3
Kf3s1k1
Kf3s1k3
Kf3s4k2
Kf3s4k4
Kf3s5k2
Kf3s6k1
Kf3s7k1
Kf3s7k3
Kf3s8k2

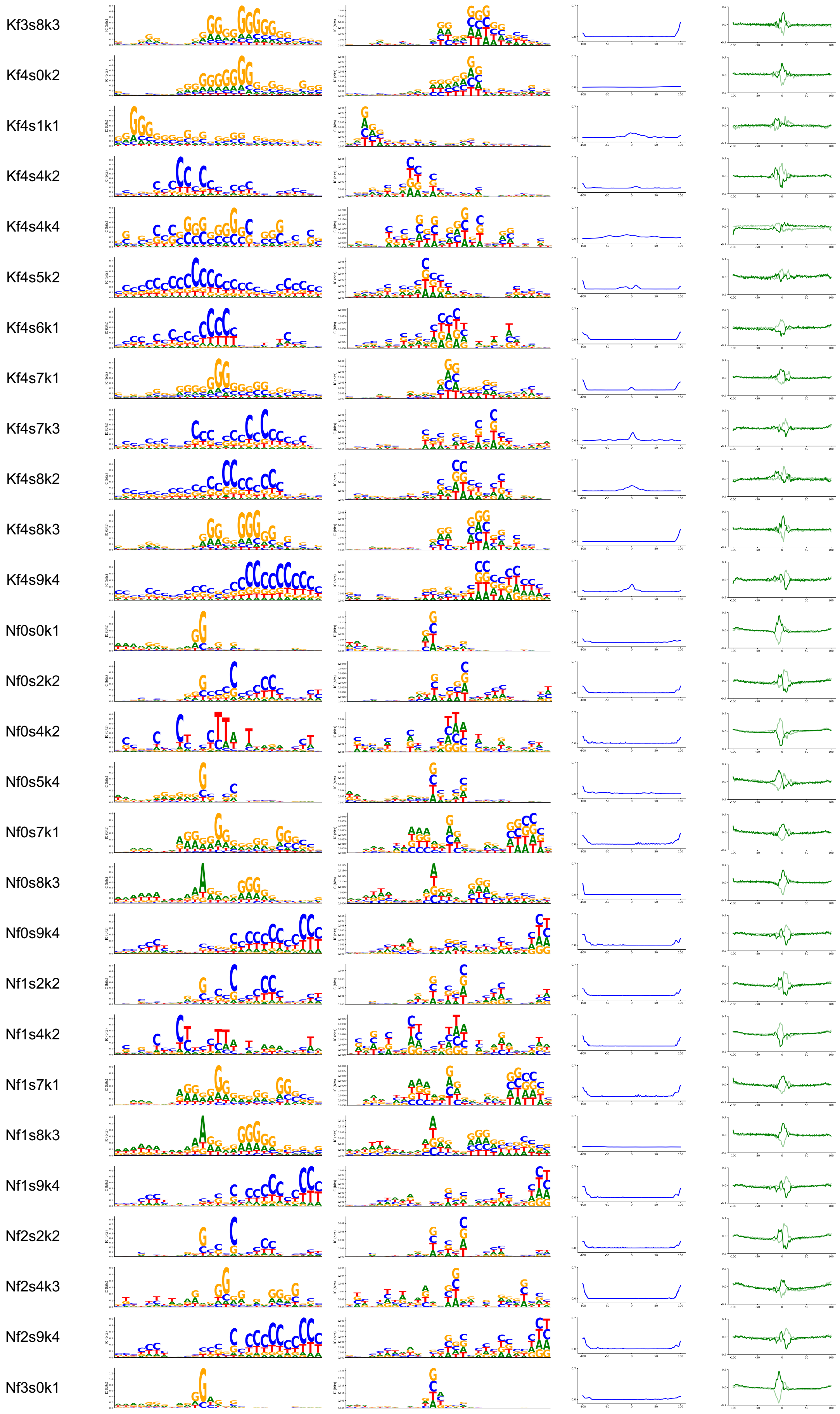

Kf3s8k3
Kf4s0k2
Kf4s1k1
Kf4s4k2
Kf4s4k4
Kf4s5k2
Kf4s6k1
Kf4s7k1
Kf4s7k3
Kf4s8k2
Kf4s8k3
Kf4s9k4
Nf0s0k1
Nf0s2k2
Nf0s4k2
Nf0s5k4
Nf0s7k1
Nf0s8k3
Nf0s9k4
Nf1s2k2
Nf1s4k2
Nf1s7k1
Nf1s8k3
Nf1s9k4
Nf2s2k2
Nf2s4k3
Nf2s9k4
Nf3s0k1

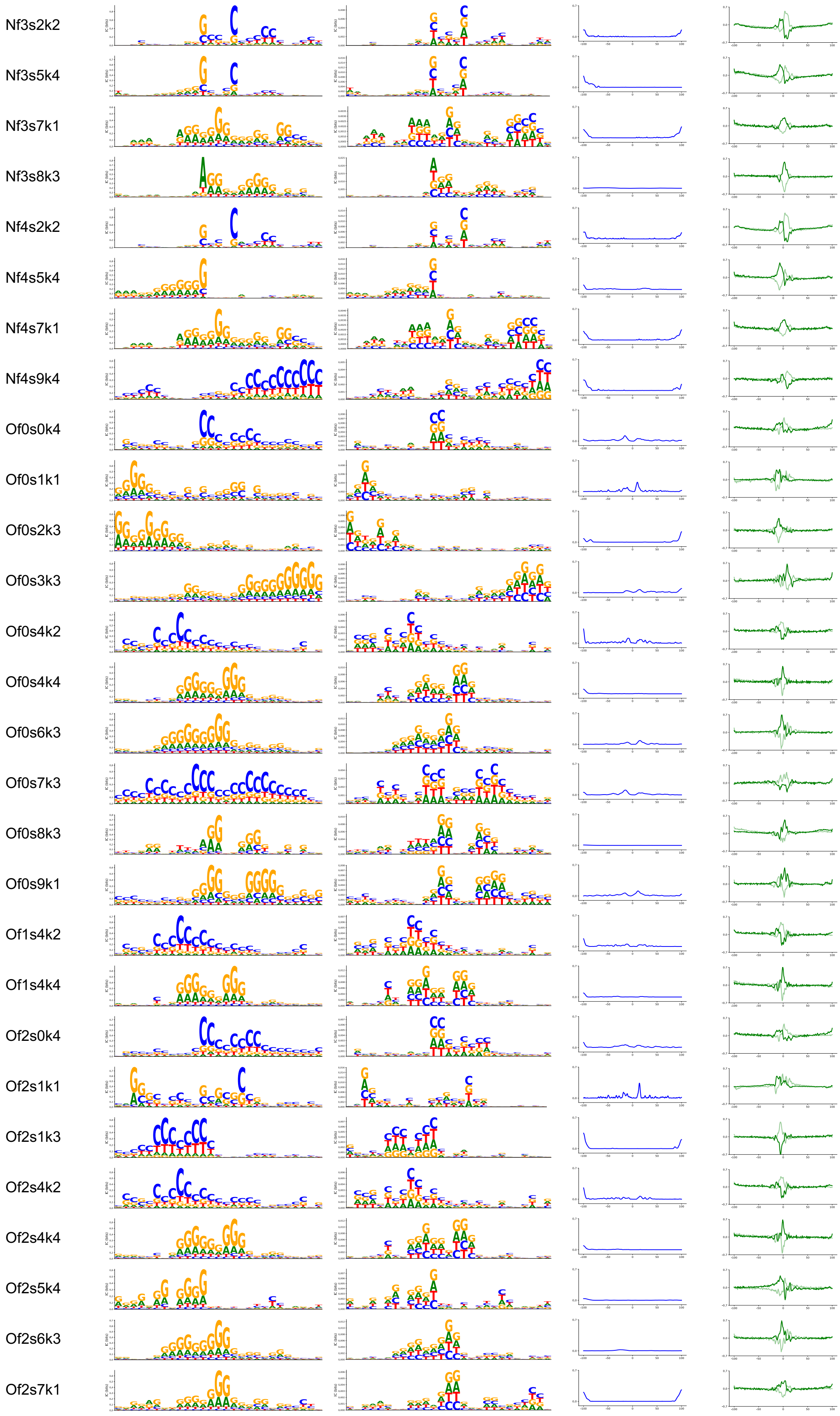
Nf3s2k2
Nf3s5k4
Nf3s7k1
Nf3s8k3
Nf4s2k2
Nf4s5k4
Nf4s7k1
Nf4s9k4
Of0s0k4
Of0s1k1
Of0s2k3
Of0s3k3
Of0s4k2
Of0s4k4
Of0s6k3
Of0s7k3
Of0s8k3
Of0s9k1
Of1s4k2
Of1s4k4
Of2s0k4
Of2s1k1
Of2s1k3
Of2s4k2
Of2s4k4
Of2s5k4
Of2s6k3
Of2s7k1

Of2s8k3
Of3s0k4
Of3s4k2
Of3s4k4
Of3s6k3
Of3s8k3
Of4s0k4
Of4s1k1
Of4s2k3
Of4s4k2
Of4s6k3
Sf0s0k3
Sf0s0k4
Sf0s1k2
Sf0s2k3
Sf0s5k1
Sf0s5k4
Sf0s6k3
Sf0s7k3
Sf0s7k4
Sf1s0k3
Sf1s1k2
Sf1s3k3
Sf1s4k2
Sf1s5k1
Sf1s5k4
Sf1s7k3
Sf1s7k4

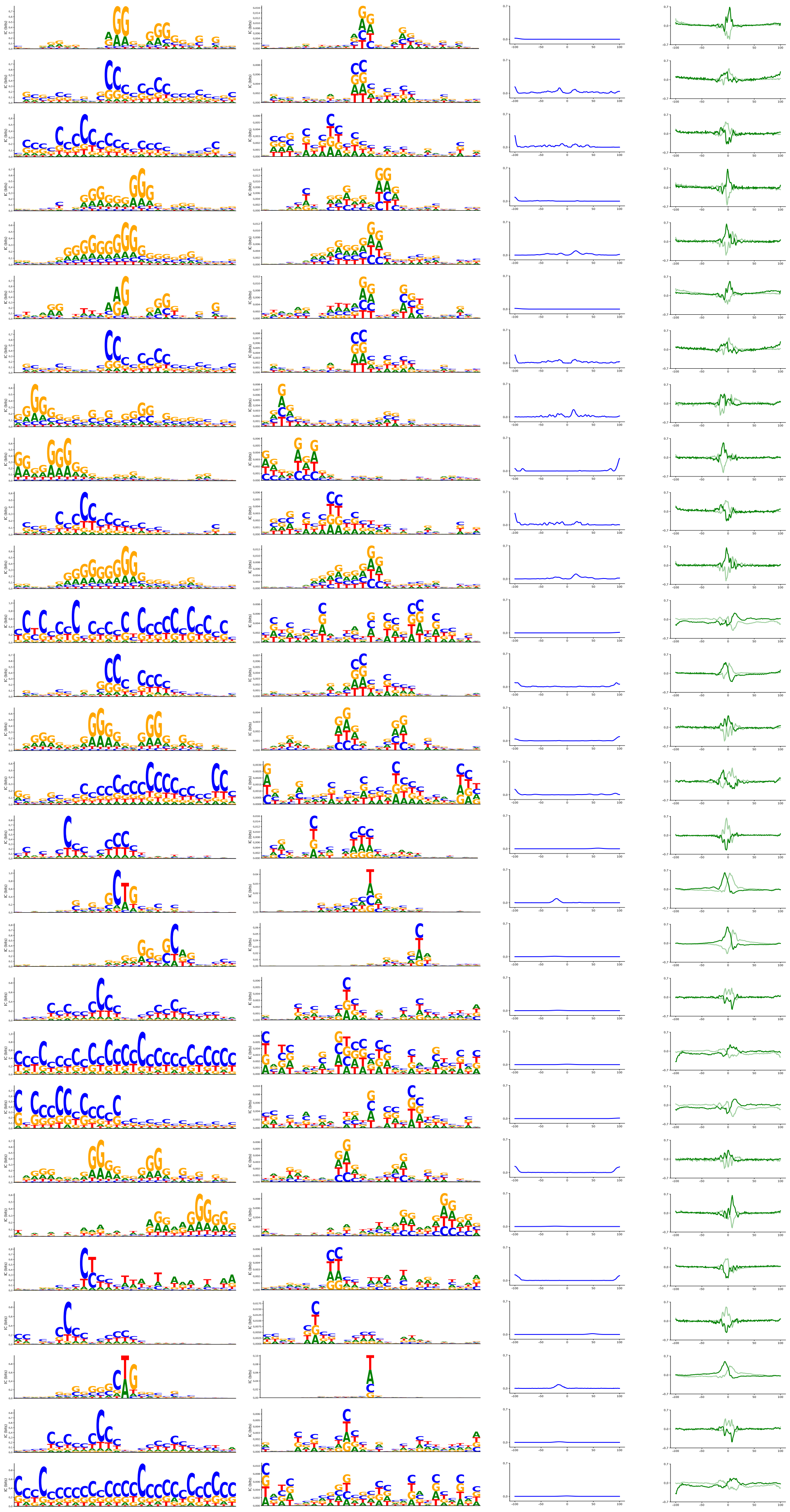

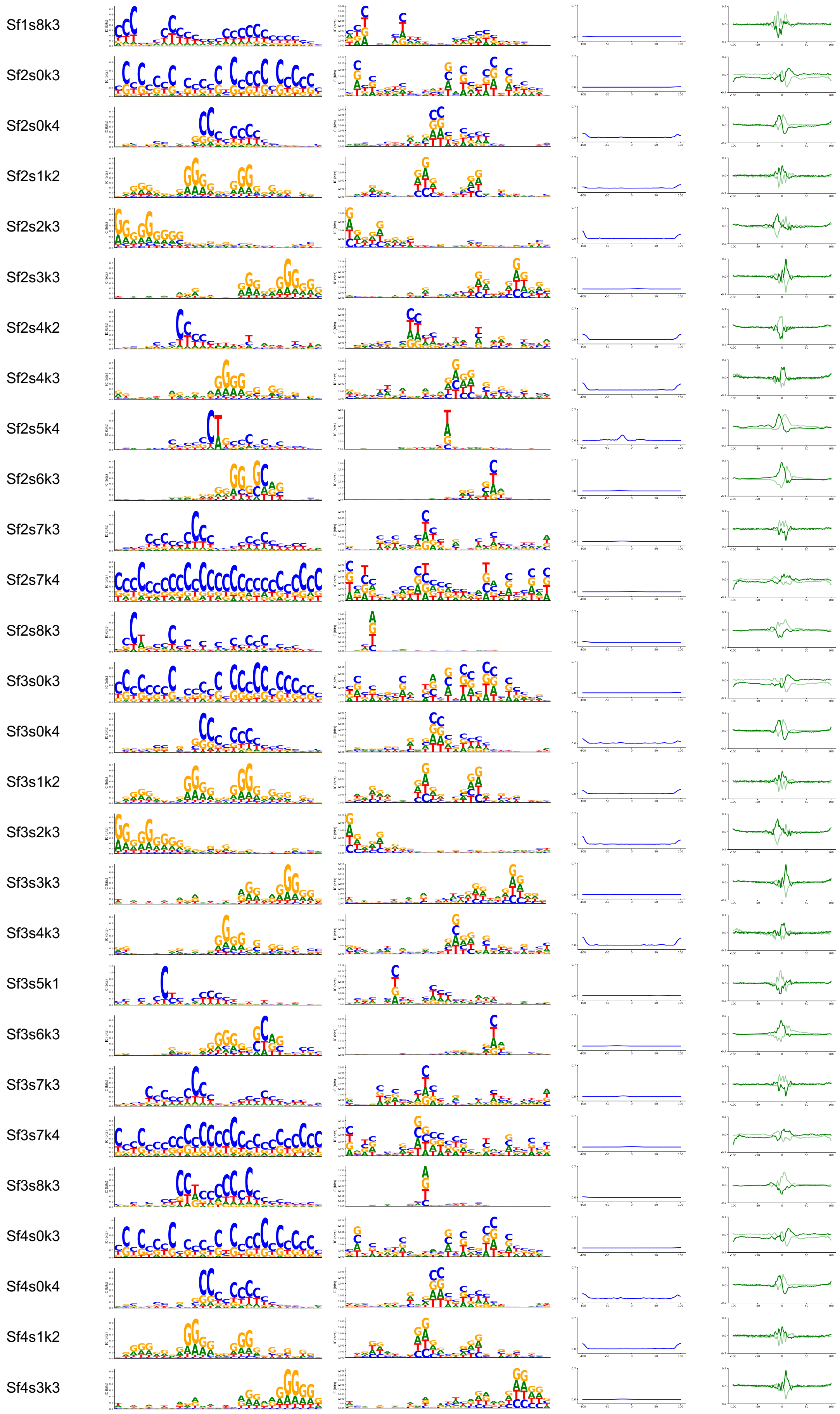
Sf1s8k3
Sf2s0k3
Sf2s0k4
Sf2s1k2
Sf2s2k3
Sf2s3k3
Sf2s4k2
Sf2s4k3
Sf2s5k4
Sf2s6k3
Sf2s7k3
Sf2s7k4
Sf2s8k3
Sf3s0k3
Sf3s0k4
Sf3s1k2
Sf3s2k3
Sf3s3k3
Sf3s4k3
Sf3s5k1
Sf3s6k3
Sf3s7k3
Sf3s7k4
Sf3s8k3
Sf4s0k3
Sf4s0k4
Sf4s1k2
Sf4s3k3

Sf4s5k4

Sf4s6k3

Sf4s7k3

Sf4s7k4

Sf4s9k4

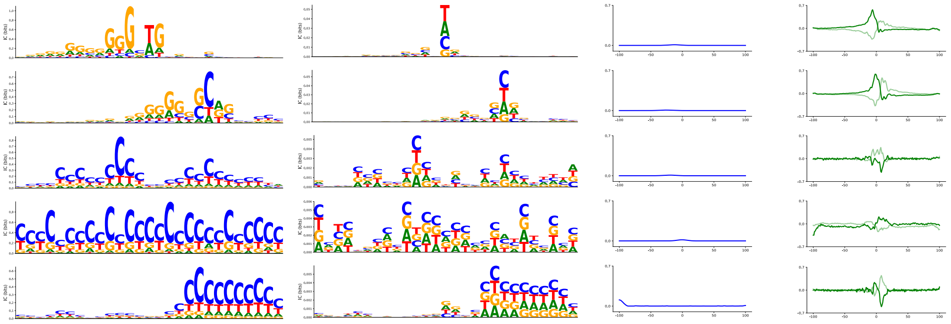

Cluster3

| ID | Seq PWM | Filter PWM | Influence Profile | Footprint Profile |
|---|---|---|---|---|

Kf1s3k1

Kf4s8k1

Nf0s1k2

Nf0s2k4

Nf0s3k1

Nf0s4k4

Nf0s6k2

Nf0s7k2

Nf1s0k1

Nf1s2k4

Nf1s3k1

Nf1s4k1

Nf1s4k4

Nf1s7k2

Nf2s0k1

Nf2s1k2

Nf2s2k4

Nf2s3k1

Nf2s4k4

Nf2s6k2

Nf2s7k2

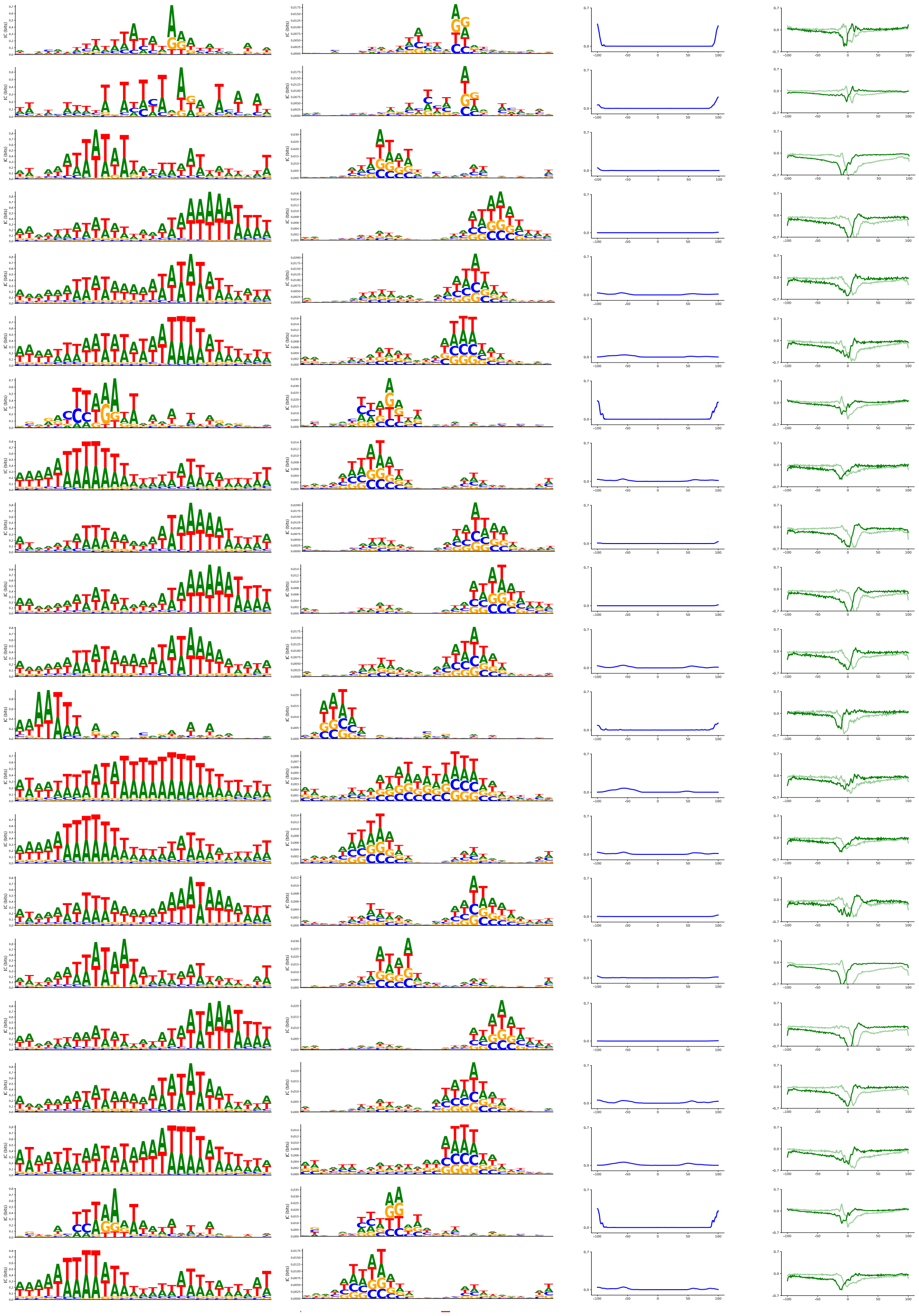

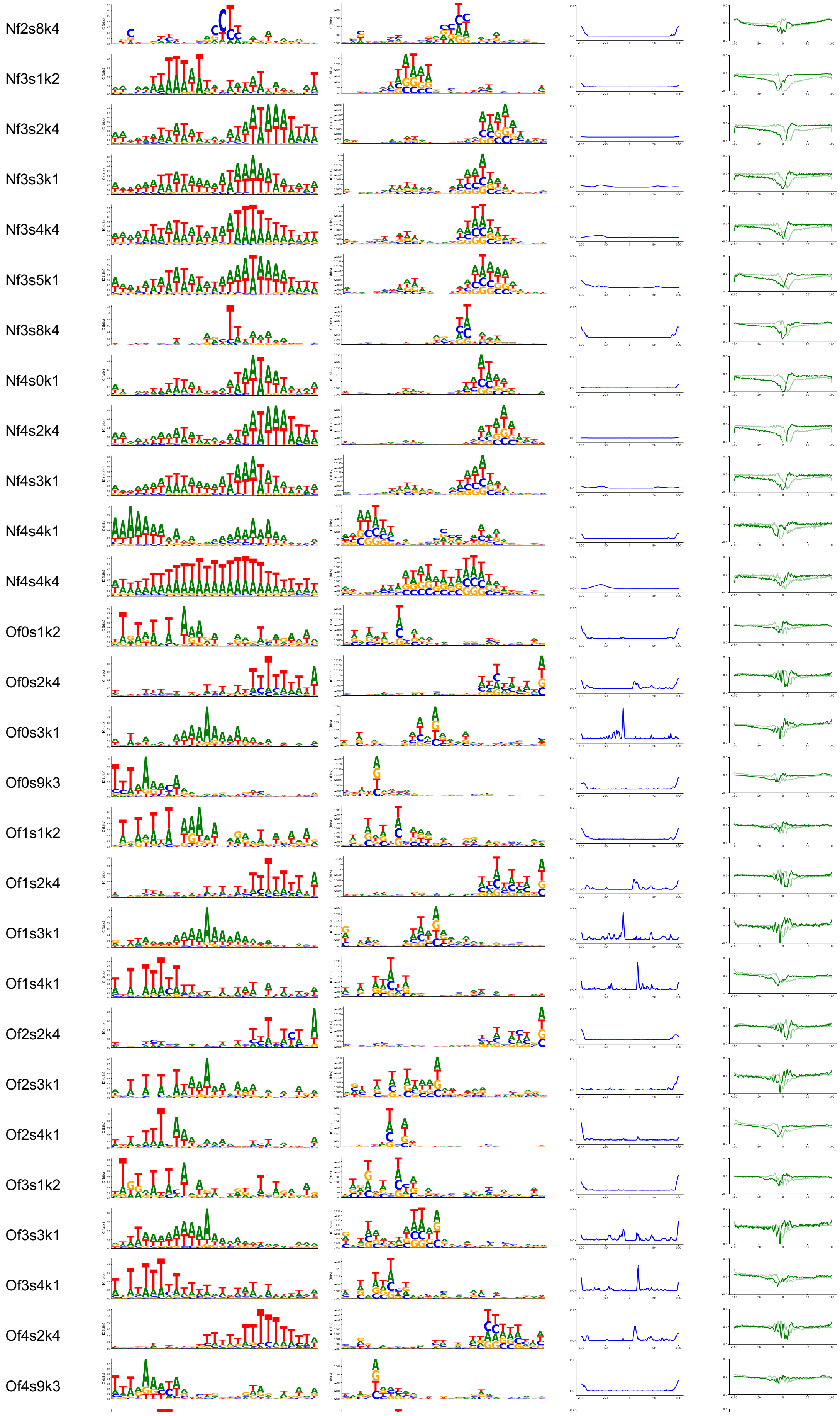
Nf2s8k4
Nf3s1k2
Nf3s2k4
Nf3s3k1
Nf3s4k4
Nf3s5k1
Nf3s8k4
Nf4s0k1
Nf4s2k4
Nf4s3k1
Nf4s4k1
Nf4s4k4
Of0s1k2
Of0s2k4
Of0s3k1
Of0s9k3
Of1s1k2
Of1s2k4
Of1s3k1
Of1s4k1
Of2s2k4
Of2s3k1
Of2s4k1
Of3s1k2
Of3s3k1
Of3s4k1
Of4s2k4
Of4s9k3

Sf0s1k4
Sf0s3k1
Sf0s9k3
Sf1s1k4
Sf1s2k4
Sf1s3k1
Sf1s9k2
Sf2s1k4
Sf2s2k4
Sf2s3k1
Sf3s1k4
Sf3s2k4
Sf3s9k3
Sf4s1k4
Sf4s3k1
Sf4s9k2

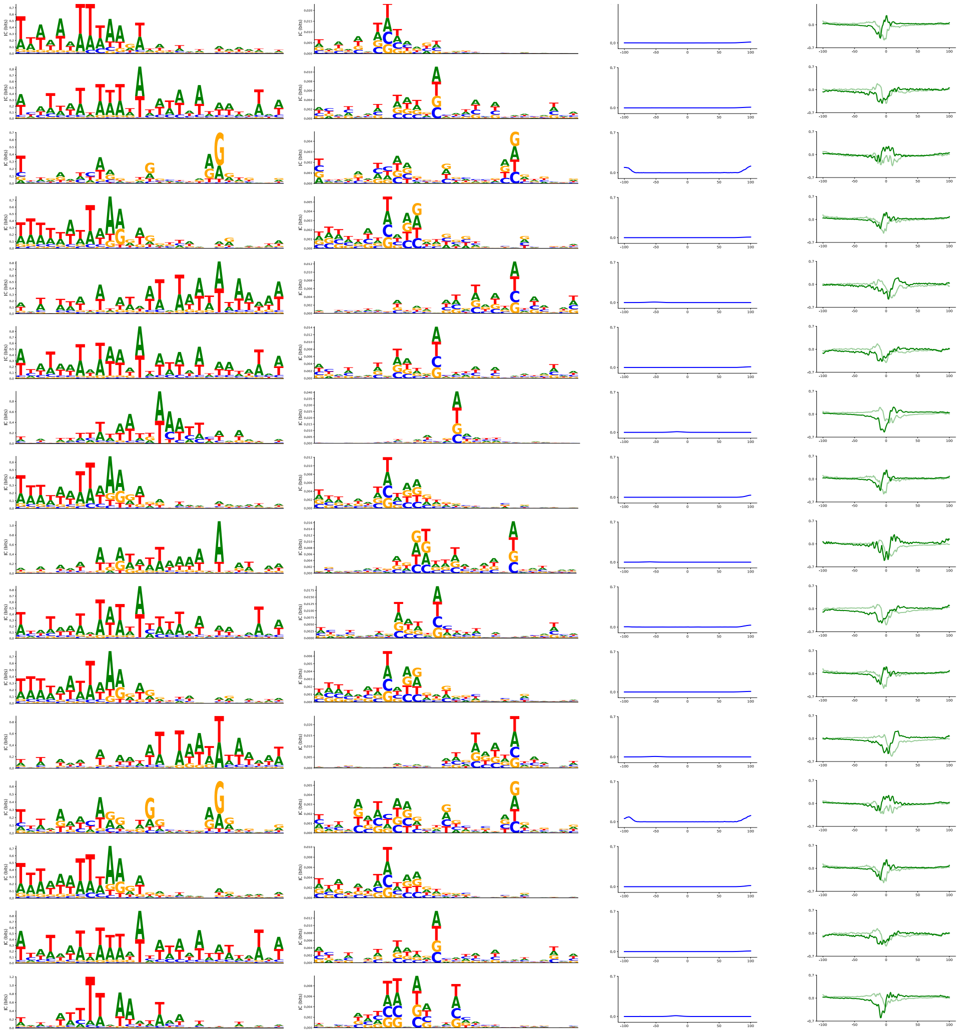

Cluster4

| ID | Seq PWM | Filter PWM | Influence Profile | Footprint Profile |
| --- | --- | --- | --- | --- |

Kf0s2k2
Kf1s2k2
Kf2s2k2
Kf3s2k2
Kf4s2k2
Of0s2k2
Of1s2k2
Of2s2k2
Of3s2k2
Of4s2k2

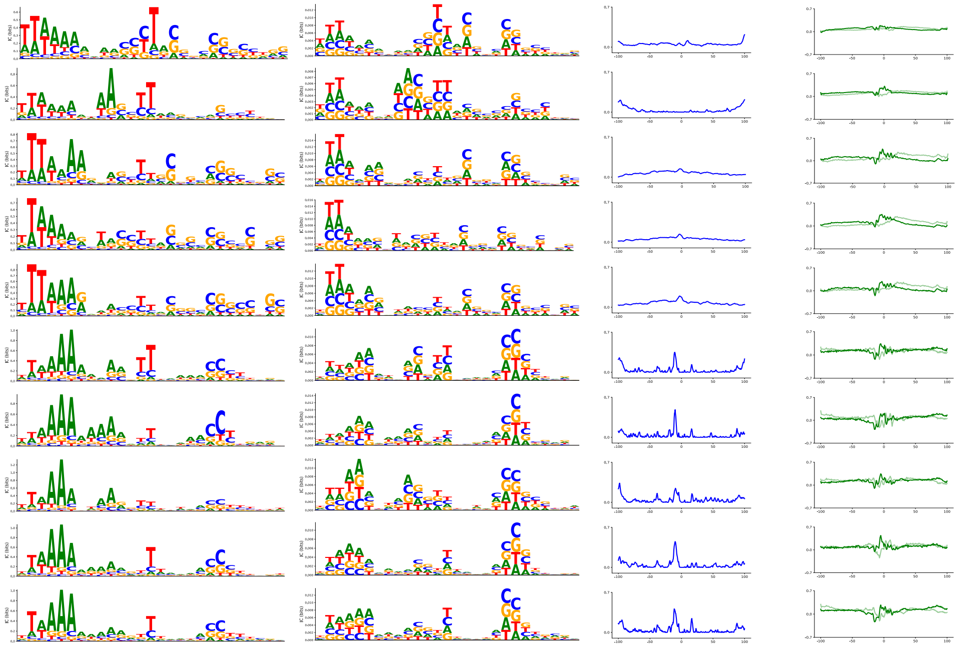

Cluster5

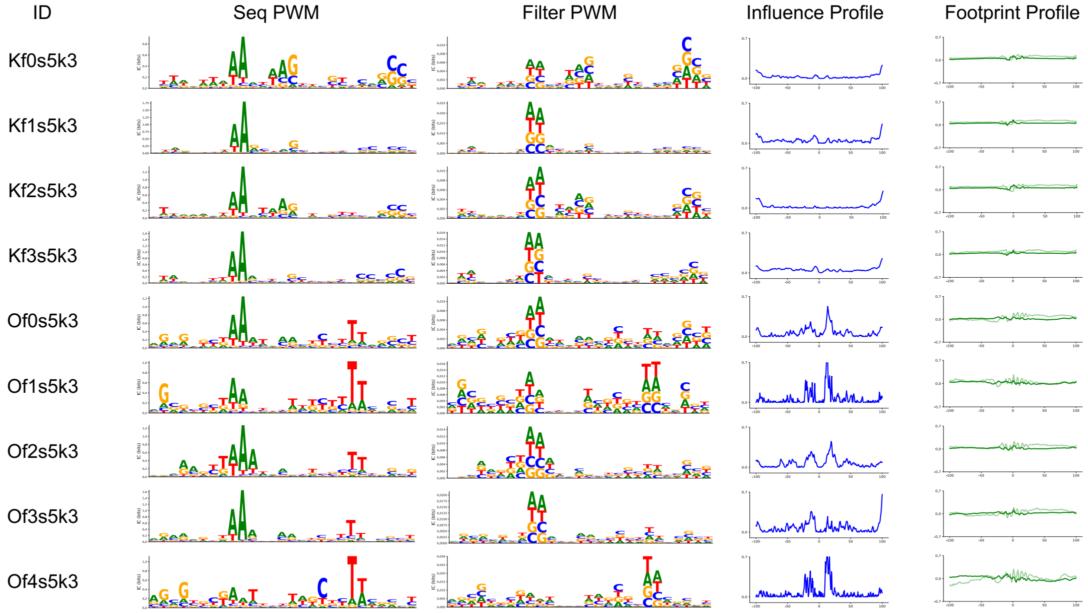


Cluster6

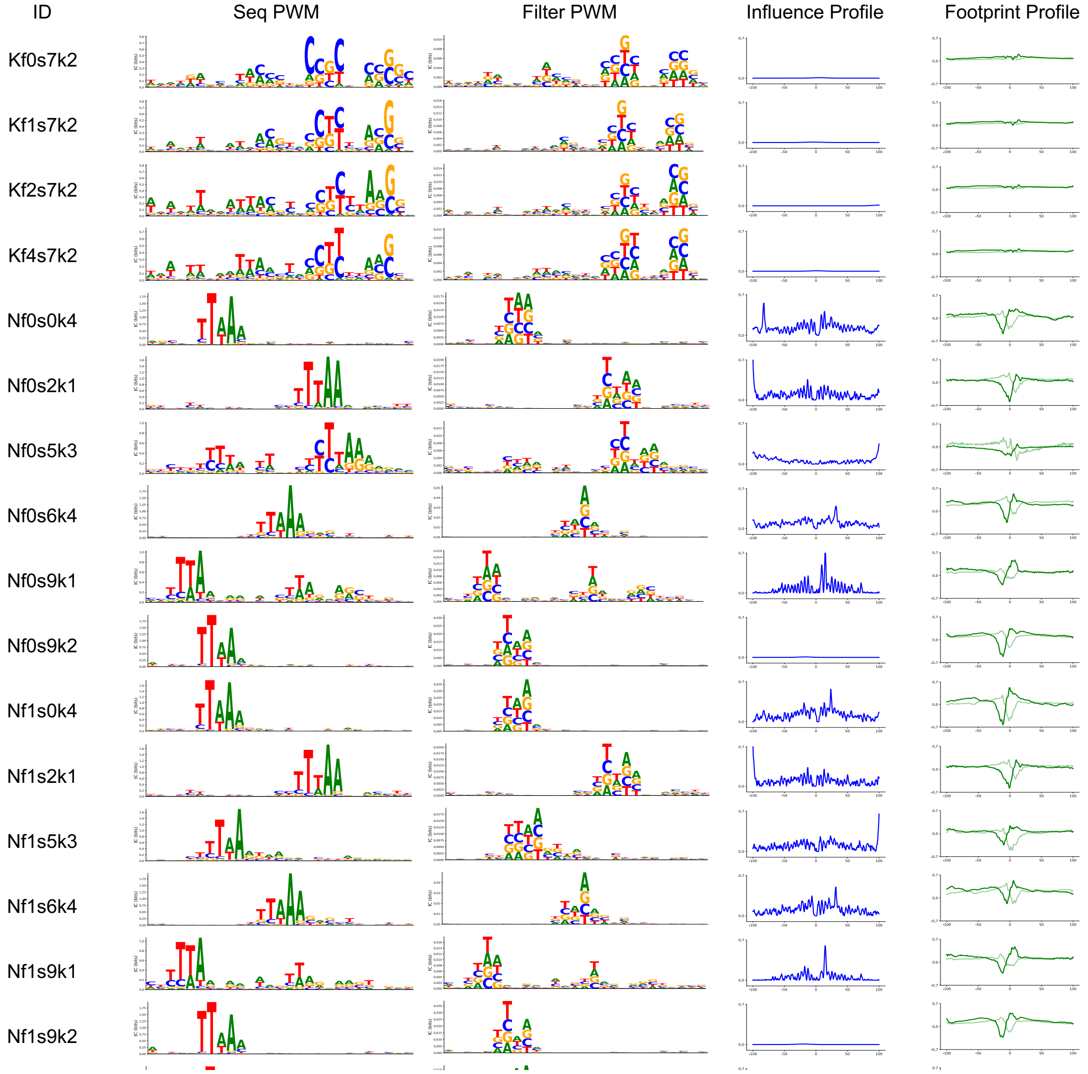

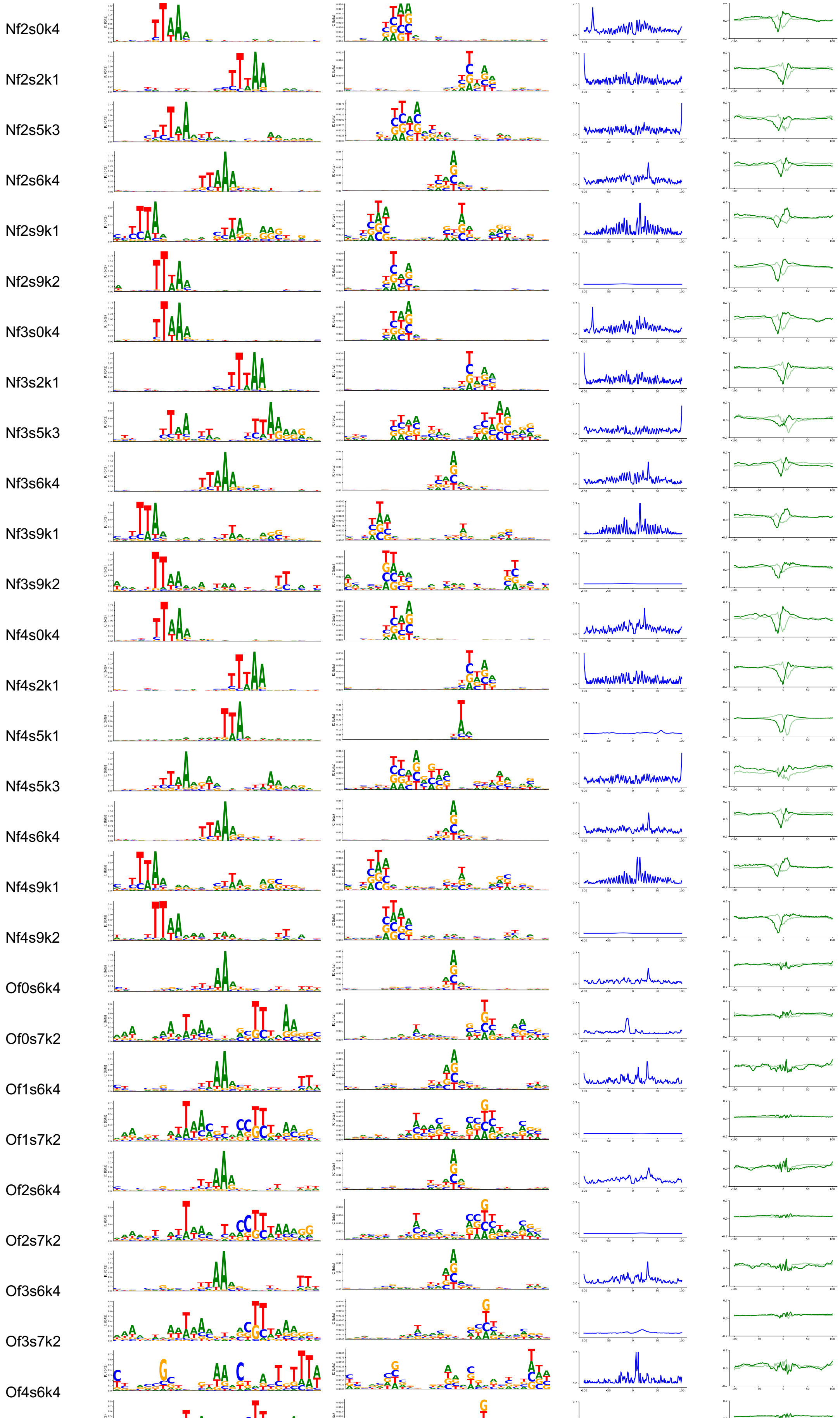
Nf2s0k4
Nf2s2k1
Nf2s5k3
Nf2s6k4
Nf2s9k1
Nf2s9k2
Nf3s0k4
Nf3s2k1
Nf3s5k3
Nf3s6k4
Nf3s9k1
Nf3s9k2
Nf4s0k4
Nf4s2k1
Nf4s5k1
Nf4s5k3
Nf4s6k4
Nf4s9k1
Nf4s9k2
Of0s6k4
Of0s7k2
Of1s6k4
Of1s7k2
Of2s6k4
Of2s7k2
Of3s6k4
Of3s7k2
Of4s6k4

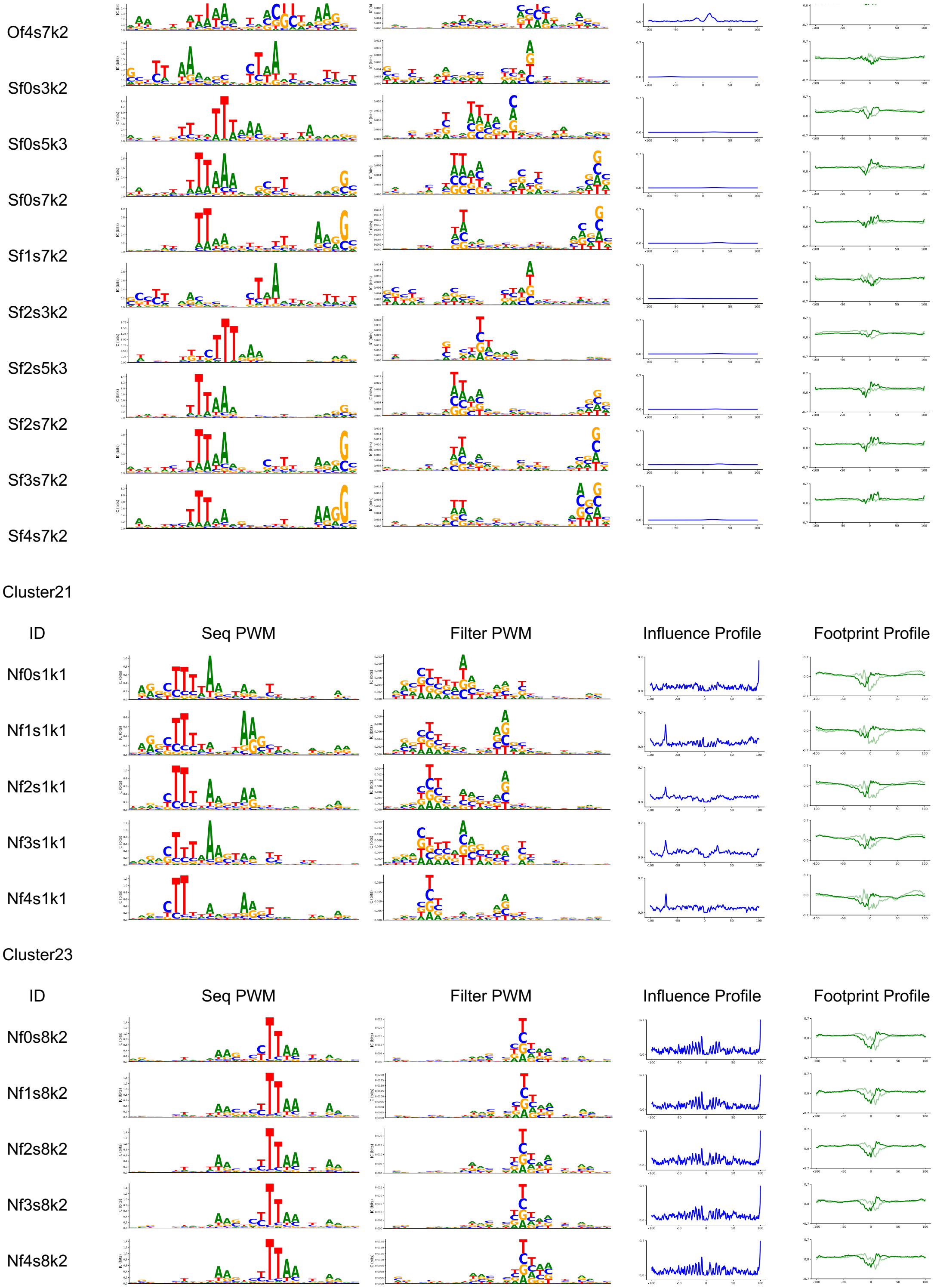
Of4s7k2
Sf0s3k2
Sf0s5k3
Sf0s7k2
Sf1s7k2
Sf2s3k2
Sf2s5k3
Sf2s7k2
Sf3s7k2
Sf4s7k2
Cluster21
ID
Seq PWM
Filter PWM
Influence Profile
Footprint Profile
Nf0s1k1
Nf1s1k1
Nf2s1k1
Nf3s1k1
Nf4s1k1
Cluster23
ID
Seq PWM
Filter PWM
Influence Profile
Footprint Profile
Nf0s8k2
Nf1s8k2
Nf2s8k2
Nf3s8k2
Nf4s8k2
